# Developing new technologies to protect ecosystems: planning with adaptive management

**DOI:** 10.1101/2024.10.24.619976

**Authors:** Luz V. Pascal, Iadine Chadès, Matthew P. Adams, Kate J. Helmstedt

## Abstract

Technology development is an essential investment for policymakers to address contemporary global crises, including climate change, biodiversity loss, the energy transition, and emergent infectious diseases. However, investing limited resources in the development of new technologies is risky. The research and development process is unpredictable, with unknown timelines and outcomes. In addition, even after successful development, the effects of deploying a new technology remain uncertain. When confronted with these uncertainties, policymakers must determine how long they should allocate resources to developing new technologies. Informed decisions require anticipating possible successes and failures of both technology development and deployment, which is a challenging optimisation task when managing dynamic systems, such as threatened ecological systems. Using an adaptive management approach from Artificial Intelligence, we discover a time limit new technologies should be developed for, which balances costs, benefits, and uncertainties during development and deployment. We extract clear and transparent general rules for investing in new technologies, building on an analytical approximation. Using Australia’s Great Barrier Reef as a case study, we demonstrate how characteristics of the managed system influence the optimal investment strategy. Our approach can inform the development of new technologies in multiple domains including biodiversity conservation, public health, energy production, and the technology industry more broadly.

**Significance:** Technology development is essential to address the crises our world faces, such as ecosystem collapse. With limited resources, policymakers must decide whether to invest in developing new technologies and, if ever, when to stop. Informed decisions require anticipating possible failures of both technology development and deployment, a challenging task when dealing with changing systems. Using an Artificial

Intelligence approach, we find a time limit for technology development that depends on characteristics of the managed ecosystem. This work can guide technology investments in many domains such as biodiversity conservation, epidemiology, energy production and the technology industry more broadly.

---

Our world is facing unprecedented crises, from biodiversity extinction [10] to climate change [6], and increased risks of infectious disease outbreaks [9]. New technologies offer an attractive opportunity to help address these challenges, e.g., carbon capture to mitigate climate change [75], genetic adaptation for species in vulnerable ecosystems [50; 5], and new vaccine development methodologies [3]. However, technology development and deployment are rife with failure and uncertainty [49], for two main reasons. First, Research & Development (R&D) is prone to the risk of failure, potentially resulting in no new technology despite development efforts [49]. Only 20% of all R&D pharmaceutical projects are successful [62]. Second, the impact a new technology will have when it is deployed is often uncertain [35; 44; 43], especially when technologies are designed for complex and dynamic systems [35]. In the face of these uncertain outcomes during development and deployment, decision-makers must decide how long limited resources should be invested on developing new technologies, and when (if ever) investments should be stopped. We address this decision problem using an adaptive management approach from Artificial Intelligence (AI). We derive general rules for investing in technology development that depend on attributes of both the technology and the system it will affect. Our approach fills a gap in the R&D planning literature by anticipating an optimal deployment strategy for new technologies that maximises expected benefits while explicitly accounting for deployment uncertainties. We apply our approach to Australia’s Great Barrier Reef, a compelling example of how new technology investments may aid in preventing or delaying the collapse of one of the world’s most iconic ecosystems [14; 50].

To plan how long we should develop a technology, we must anticipate two things: the success or failure of its development, and the benefits it will bring if and when it is deployed in the field. To date, the R&D planning literature suffers from a lack of approaches to anticipate possible successes and failures of both technology development and deployment. While the uncertain success of technology development is often accounted for in R&D decision-making models [27; 25; 2; 36; 26; 54], these decision tools assume that the development process follows a known model structure (e.g. [26] uses a Poisson process to describe technological advancements). However, in practice, uncertainty in that development model structure can lead to sub-optimal or risky decisions. Even after accounting for development uncertainty, underestimating uncertainties of deployment outcomes can result in harmful unintended consequences. For example, circle hooks, initially designed to reduce sea turtle mortality in fisheries, resulted in an increased capture of threatened sharks [43]. R&D literature provides little guidance on how to anticipate the uncertain outcomes of technology deployment, especially when dealing with dynamic systems [16; 46; 64; 2; 36]. Two broad approaches are used to anticipate the benefits of deploying a new technology. One of these approaches, prone to human errors and biases [35; 42], is to elicit uncertainty about future deployment outcomes from expert knowledge before development [22; 25; 2; 36]. The other, more technical approach is to fully resolve uncertainties about deployment outcomes through an iterative process that alternates technology development and small-scale sequential experimentation [16; 64]. However, this technical approach assumes that deployment outcomes are deterministic after experimentation. This assumption is reasonable in many domains, but technology deployment uncertainties can rarely be fully resolved when dealing with complex dynamic systems [35; 42].

Here, we show how adaptive management can optimally guide investments in technology development and deployment in the same planning process. Adaptive management is the recommended best practice in ecology and natural resource management to manage dynamic systems under uncertainty [45]. Solving an adaptive management problem means finding a strategy to maximise expected benefits over time while accounting for the uncertain system’s response to management actions [29; 65; 51; 20; 58]. Adaptive management can anticipate and account for uncertain outcomes so is ideally suited for planning technology development and deployment. Yet this application remains unexplored. We frame both technology development and deployment as two sequential adaptive management problems that we solve as a Partially Observable Markov Decision Processes (POMDPs) [7; 19; 29; 61] (see Methods). The first adaptive management problem optimally allocates resources to developing a technology with uncertainty in the model that predicts development success (see Methods). The solution of this first problem depends on the future expected benefits achieved by technology deployment. Therefore, to reach optimality, we calculate the maximum expected deployment outcomes with a second adaptive management problem that optimally plans technology deployment, this time with uncertainty in the model that predicts deployment success (see Methods).

We illustrate our approach using an example of planning technology development and deployment for Australia’s Great Barrier Reef. The Great Barrier Reef is the richest coral reef ecosystem in the world [31], and significantly contributes to the Australian economy by providing important ecosystem services [23; 39; 5]. However, due to growing pressures such as climate-related events [33; 37; 68], poor water quality [74] and outbreaks of coral predators [63], the development of new technologies is a clear option to supplement existing management practices, and avoid ecosystem collapse [4; 5; 55] with over $300 million AUD (over $200 million USD) already invested in technology development and deployment [50]. Examples of new technologies under development include water cooling and shading techniques to reduce heat stress, three-dimensional physical structures to increase reef recovery, and biological control of coral predator impacts using genetic engineering approaches [11]. According to a recent Great Barrier Reef R&D report [50], the average costs of technology development and deployment are $5 million AUD ($3.2 million USD) and $10 million AUD ($6.4 million USD) respectively, per technology and per year.

We model the development of a new technology for managing the Great Barrier Reef, as a two-state stochastic dynamic system that can only transition from *idle* to *ready* (see Methods and SI A). Given the uncertainty surrounding future development success, the decision-maker begins with an initial belief that the technology can be successfully developed (*b*_0_, see Methods and SI A). This belief is updated as the decision-maker receives annual feedback about the development’s success (see eq. 1). The belief decreases until it reaches the threshold for stopping development or takes 1 if the technology is successfully developed (see Methods and SI C). The optimal adaptive management strategy for technology development provides R&D investment recommendations based on the current belief in the technology’s successful development (*b*_*t*_), the cost of technology development (*C*_dev_), and expected deployment outcomes (*R*_dep_). To calculate the maximum expected outcomes of technology deployment (maximum achievable *R*_dep_), we model the Great Barrier Reef ecosystem as a two-state stochastic dynamic system that can transition between *healthy* and *unhealthy* states (second adaptive management problem, see Methods and SI B). When *healthy*, we assume that the ecosystem produces $6.4 billion AUD ($4.3 billion USD) in ecosystem services annually [59], but when unhealthy, we estimate that the ecosystem services produced drop to 67% of this yearly value (see SI B.1). We use long-term monitoring data from the Australian Institute of Marine Science [8; 1] to estimate the ecosystem dynamics under current management: the degradation probability is estimated at 0.8 (*p*_d_: yearly probability of degrading from healthy to unhealthy) and the recovery probability at 0.2 (*p*_r_: yearly probability of recovering from unhealthy to healthy) (Methods and SI B.1). The adaptive management strategy for technology deployment provides yearly recommendations (at year *t*) as a function of the beliefs that the technology is beneficial for restoration (improving transition from unhealthy to healthy, 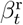) and will prevent decline (avoiding transition from healthy to unhealthy, 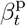) (see Methods and SI B.2). We examine how the technology development strategy is also influenced by these ecosystem characteristics: the beliefs that the technology is beneficial for the ecosystem (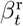 and 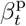), and the natural probabilities of decline and recovery of the ecosystem (*p*_d_ and *p*_r_). Finally, we provide general insights on the technology development and deployment strategies for systems with different degradation and recovery probabilities.

## Results

### Technology development

It is optimal to invest in technology development for at most *T*_max_ years, and stop investments if the technology is not successfully developed during that *T*_max_-year period (Fig. 1). This value of *T*_max_ can be unambiguously determined as an outcome of solving the technology development POMDP (first adaptive management problem, see Methods and SI A). During the development phase, the decision maker receives a yearly baseline benefit (*R*_BAU_, corresponding to the system’s services generated under business-as-usual activities, see Methods and SI A) and pays the ongoing technology development cost (*C*_dev_ per year). The technology may be successfully developed during that *T*_max_-year period, which leads the decision-maker to deploy the technology and perceive the yearly benefits that the technology generates (*R*_dep_). Conversely, if R&D investments repeatedly fail to develop a new technology, the optimal time to stop development investments (*T*_max_) depends on the belief that the technology will be successfully developed (*b*_*t*_), and on how much the expected deployment benefits (*R*_dep_) exceeds the balance of baseline benefit (*R*_BAU_) and development costs (*C*_dev_); that is *T*_max_ depends on Δ_*R*_ = *R*_dep_ − (*R*_BAU_ − *C*_dev_). This dependence is supported by our analytical approximation (dashed lines in Fig. 2, see also Eq. 2 and 3), demonstrating the importance of the anticipated expected deployment benefits when planning technology development. A higher belief in technology development success (*b*_*t*_) leads to a longer time limit for technology development (*T*_max_, see Fig. S6). Expensive technologies require a higher belief in technology development success to offset development investments (Fig. S6). For the Great Barrier Reef case study (see Table S1 for parameters), when the belief in technology development success is 0.5, it is optimal to invest in technology development for around 40 years before surrendering; this drops to 20 years if the belief is 0.1 (Fig. S6).

**Figure 1.**
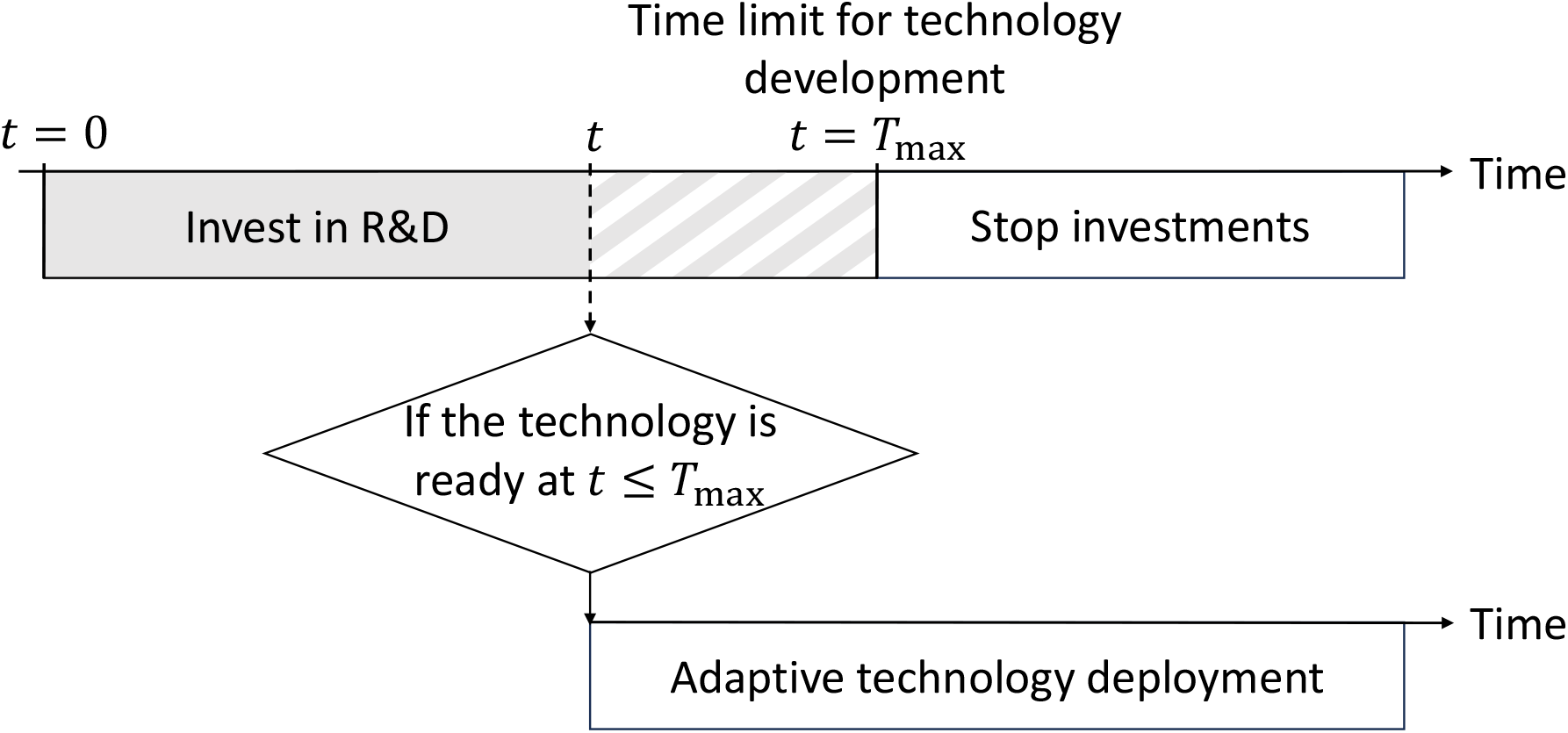
Timeline of the optimal investment strategy. Decisions are made at each time step (along the horizontal line). It is optimal to invest in project development for at most *T*_max_ years. If the technology is ready at *t* ≤ *T*_max_, it is then optimal to divert development investments to deploy the new technology through an adaptive management program (striped area). However, if the time limit for technology development is reached before a technology is ready, it is optimal to stop technology development investments. The special case where *T*_max_ = 0 corresponds to the strategy where it is better to never invest in technology development.

**Figure 2.**
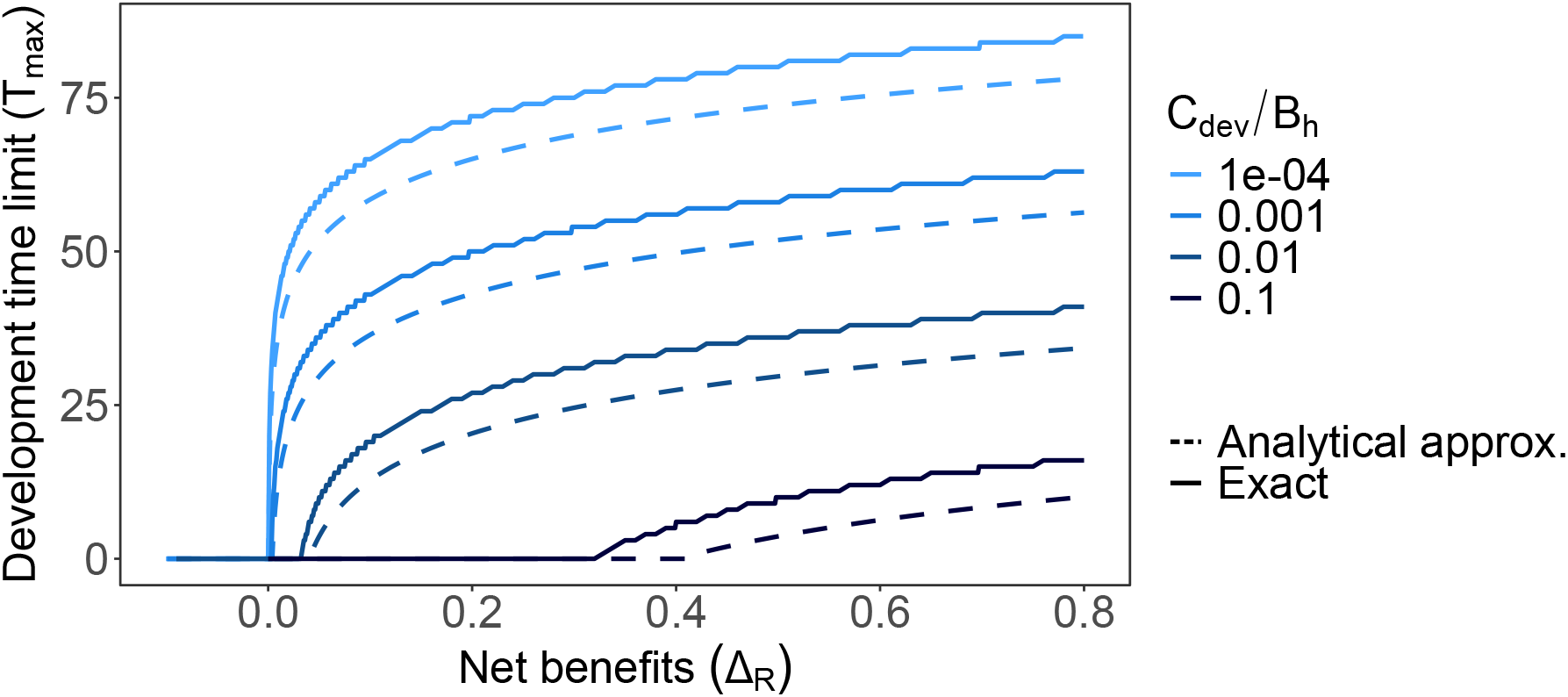
Influence on the time limit for technology development (*T*_max_) of the net benefit of technology deployment (Δ_*R*_) defined as the difference between expected outcomes of deployment (*R*_dep_) and benefits without the technology reduced by development costs (*R*_BAU_ − *C*_dev_). Dashed lines represent analytical approximations (see eq. 3). Each color depicts different costs of development *C*_dev_. Note here that all values of benefit and cost parameters (Δ_*R*_ and *C*_dev_) are scaled to the maximum achievable value of the ecosystem when healthy (*B*_h_). Here, the belief in successful technology development is 0.5.

Higher net increase of benefits due to technology deployment (Δ_*R*_) results in longer time limits for technology development (*T*_max_, Fig. 2). However, as development costs increase, the net benefits required to justify development also increase (larger values of Δ_*R*_ to obtain *T*_max_ *>* 0, Fig. 2).

### Technology deployment

Since the benefits gained when deploying a technology determine the time limit for its development (Fig. 2), we use adaptive management to find an optimal deployment strategy that maximises the expected benefits of deployment while accounting for the uncertainty about how the system will respond to the technology (maximum achievable *R*_dep_, denoted *R*_AM_). In practice, the optimal adaptive deployment strategy provides yearly recommendations about whether to deploy the technology or not. We find optimal adaptive strategies take three simple forms: never deploy, only deploy based on the system state (healthy or unhealthy), or deploy regardless of the ecosystem state (Fig. 3A). The optimal strategy depends on two beliefs the decision-maker has about the technology: that technology deployment will be beneficial for restoration 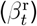 and preventing decline 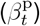. These beliefs are updated over time depending on how the system responds to the implementation of a management action (see eq. 1). For the Great Barrier Reef case study, when the initial beliefs 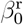 and 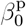 are both set to 0.8 (i.e., the decision-maker is 80% confident the technology will recover an unhealthy reef, and 80% confident it will maintain a healthy reef) and the ecosystem is initially *unhealthy*, it is optimal to implement the action *deploy*. If the ecosystem stays *unhealthy* after 6 deployment attempts, it is optimal to stop deploying the technology when the ecosystem is *unhealthy*. The beliefs update each year upon observing the system response, in a trajectory represented by black squares in Fig. 3A (other belief trajectories will occur if the initially unhealthy ecosystem responds more favourably to technology deployment).

**Figure 3.**
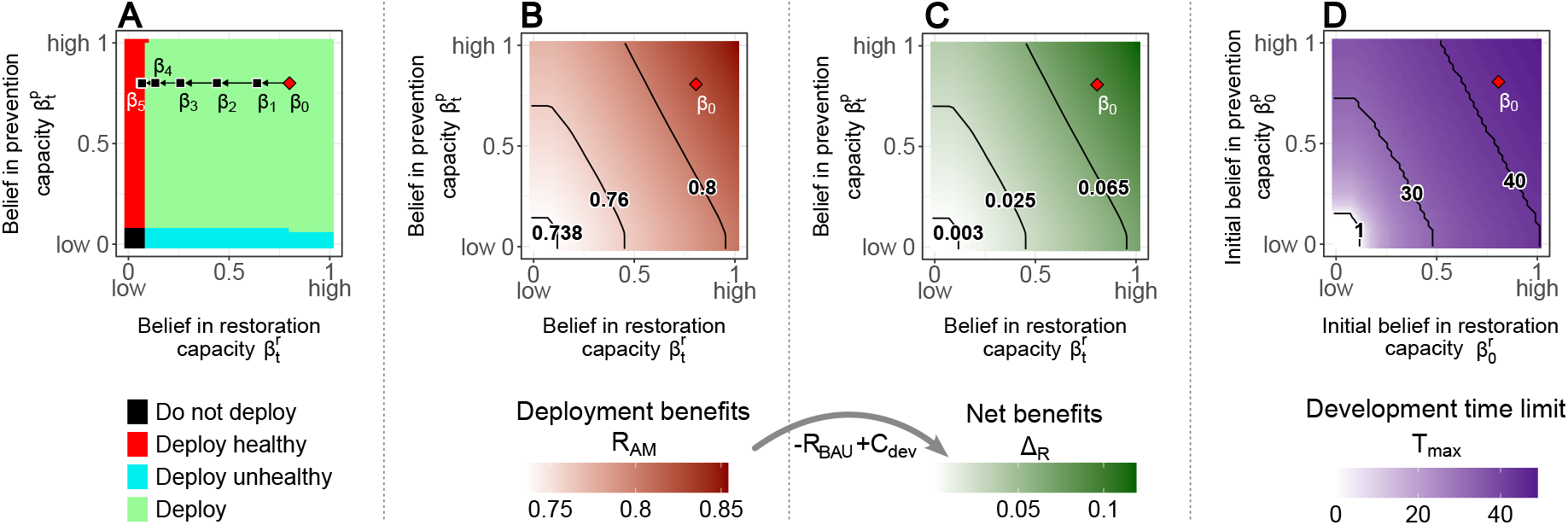
Results for the Great Barrier Reef case study. **A** Adaptive management strategy for technology deployment (independent of time). At each time step, the optimal decision is determined by the combination of beliefs that technology deployment is beneficial for ecosystem restoration or preventing decline (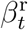 and 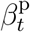). The red diamond represents the initial belief that technology deployment is beneficial regardless of current ecosystem health (*β*_0_), that can be summarized by a combination of beliefs 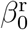 and 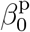 (*x* – and *y* -coordinates of the point *β*_0_ in this plot). The successive black squares represent an example of belief trajectory, after implementing the optimal action, and observing that the ecosystem is *unhealthy*. After 6 deployment attempts and no improvement to reef health, the belief that the technology is beneficial has moved to *β*_5_, indicating no further technology deployment should be attempted for this unhealthy reef. **B** Influence of the beliefs that technology deployment is beneficial for ecosystem restoration or preventing decline (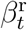 and 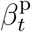) on the yearly expected benefits of technology deployment with the optimal adaptive management strategy (*R*_AM_). Darker shades indicate higher expected benefits of deployment. The black lines represent the isocurves for different possible values of *R*_AM_. **C** Influence of the beliefs that technology deployment is beneficial for ecosystem restoration or preventing decline (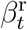 and 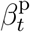) on net expected benefits (Δ_*R*_ = *R*_AM_ − *R*_BAU_ + *C*_dev_). For a given ecosystem, the value of the ecosystem services generated without the technology (*R*_BAU_) and costs of technology development (*C*_dev_) are constant. Darker shades of green illustrate higher values of Δ_*R*_. The black lines represent the isocurves for different possible values of Δ_*R*_. **D** Influence of the initial beliefs that technology deployment is beneficial for ecosystem restoration or preventing decline (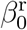 and 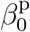) on the time limit for technology development *T*_max_. Darker shades of purple illustrate higher values of *T*_max_. The black lines represent the isocurves for different possible values of *T*_max_. Below the *T*_max_ = 1 isocurve, it is optimal to never invest in technology development (*T*_max_ = 0).

Interestingly, we identify four patterns of influence of beliefs on expected deployment outcomes (*R*_AM_ in Fig. 3B). When the technology is believed to both restore and prevent decline of the system (beliefs are high, 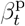 and 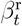 close to 1), expected deployment benefits are high (darker shades in Fig. 3B) and depend on both beliefs (diagonal isocurves in Fig. 3B). Conversely, when the technology is believed to not restore or prevent decline of the system (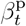 and 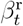 closer to 0), the expected deployment outcomes are equivalent to the expected outcomes without the technology, as the technology will never be deployed (lighter shades in Fig. 3B). If the technology is believed to restore the ecosystem but not to prevent decline (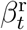 close to 1 and 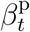 close to 0), expected deployment benefits primarily increase with the belief that the technology is effective for restoration (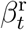 increases, vertical isocurves of *R*_AM_ in Fig. 3B). We obtain symmetric results if the technology is believed to prevent ecosystem decline but not to restore the ecosystem (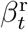 close to 0 and 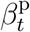 close to 1, horizontal isocurves in Fig. 3B).

Similarly to the expected deployment benefits (*R*_AM_, Fig. 3B), we observe the same four patterns of influence of beliefs (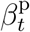 and 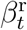) on the net benefits due to deployment (Δ_*R*_, Fig. 3C), and of the initial beliefs (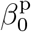 and 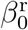) on the time limit for technology development (*T*_max_, Fig. 3D). This is because net benefits (Δ_*R*_) and the time limit (*T*_max_) depend on the expected deployment benefits (*R*_AM_, as established in the results for technology development) and because expected outcomes without the technology (*R*_BAU_) and costs of development (*C*_dev_) are both independent of the initial beliefs (see SI B.1).

The results shown for Fig. 3 assume a system with characteristics of fast degradation (*p*_d_ = 0.8) and slow recovery (*p*_r_ = 0.2) in absence of technological intervention. This reflects the Great Barrier Reef case study (see SI B.1). For systems with other degradation and recovery characteristics, the optimal deployment strategies (as in Fig. 3A) will change (Fig. 4 top row), and in turn the optimal time to stop technology development (as in Fig. 3D) will also change (Fig. 4 bottom row). For systems with fast degradation and fast recovery (e.g. North American Lodgepole Pine forests, *Pinus contorta var. latifolia*, which are sensitive to fires but regrow quickly, Fig. 4A), it is optimal to deploy the technology only when the system is healthy for most of the possible beliefs in the technology’s benefits (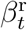 and 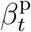 in red region in Fig. 4A(i)). For systems with these characteristics, the initial belief that the technology is beneficial for preventing decline 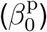 has the strongest influence on the time we can afford developing a technology (horizontal isocurves of *T*_max_ in Fig. 4A(ii)). Hence, technologies with higher confidence in preventing degradation are more likely to be beneficial for fast degrading and fast recovering systems. Systems defined by fast degradation and slow recovery (e.g. the Great Barrier Reef, Fig. 4B) are highly threatened systems. Here the dominant deployment strategy is to deploy for most beliefs (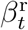 and 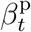 in green region in Fig. 4B(i)). This indicates that both initial beliefs (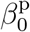 and 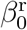) influence the time limit (diagonal isocurves of *T*_max_ in Fig. 4B(ii)). This result suggests that technologies improving restoration and/or reducing degradation are beneficial for these threatened systems. For systems with slow degradation and fast recovery (e.g. Jamaican *Alchornea latifolia*, a forest tree with high resistance to fire and high recovery response, Fig. 4C), the dominant deployment strategy is to not deploy (black region in Fig. 4C(i)). As a result, the initial beliefs in the benefits of deployment (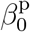 and 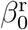) have little effect on the number of years of technology development before stopping: it is optimal to not start technology development at all (*T*_max_ = 0 in Fig. 4C(ii)). Finally, for systems with a slow degradation and slow recovery (e.g. sections of the Great Barrier Reef that are subject to less frequent but more intense cyclones, Fig. 4D) the initial belief that the technology is beneficial for restoration 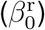 has the strongest effect on the time limit for technology development (vertical isocurves of *T*_max_ in Fig. 4D(ii)), as the dominant strategy is to deploy when the ecosystem is unhealthy (blue region in Fig. 4D(i)). Here, technologies designed for system recovery are more likely to be beneficial and worth developing. We further generalize our results to all possible systems in Fig. S7.

**Figure 4.**
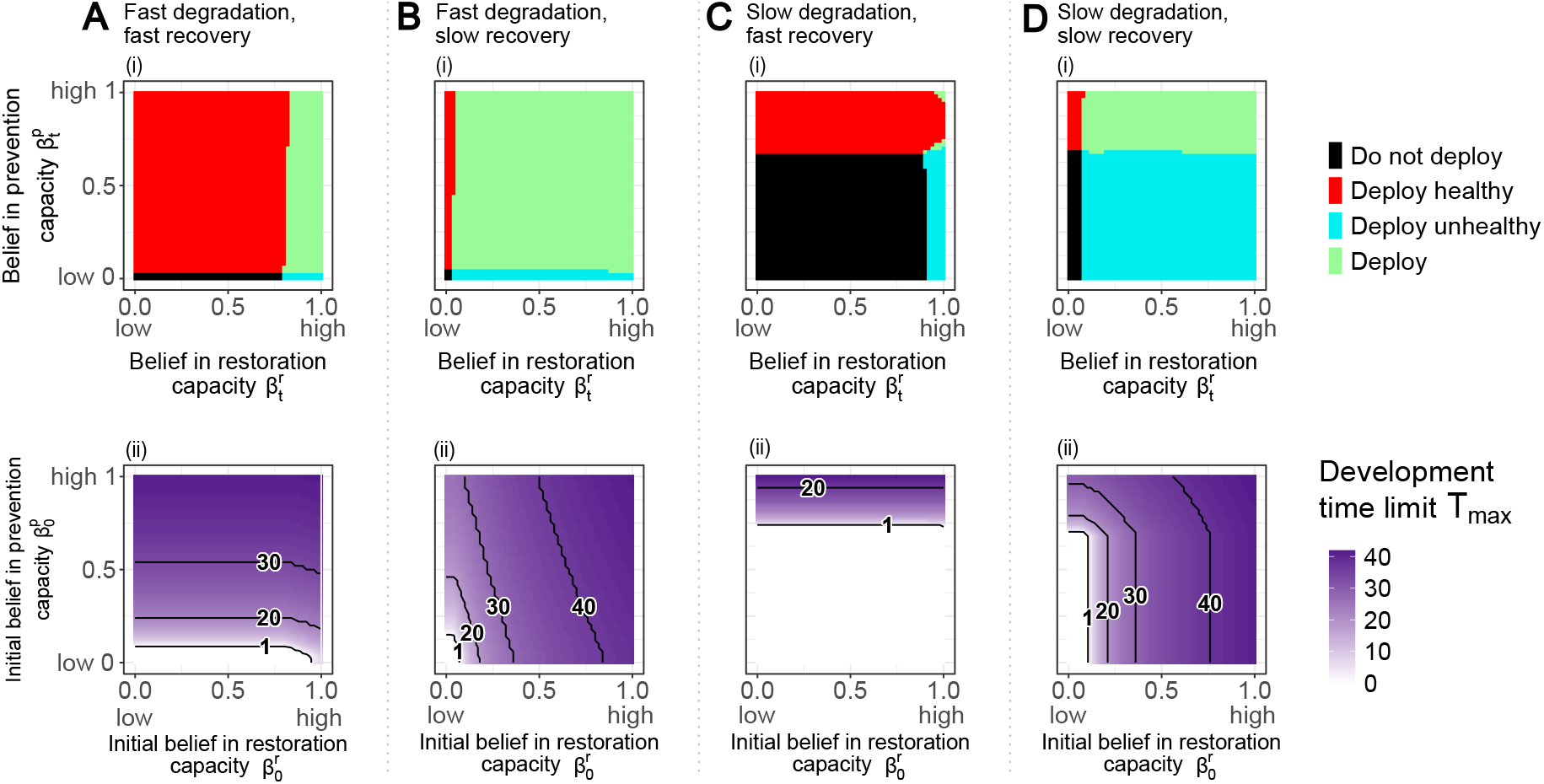
Influence of current belief in benefits of technology deployment on adaptive technology deployment strategy (top row (i)) and of the initial beliefs on the maximum number of years investing in technology development (bottom row (ii)) for four degradation-recovery profiles of ecosystems. **A** Fast degradation and fast recovery profile: *p*_d_ = 0.9, *p*_r_ = 0.9. **B** Fast degradation and slow recovery profile: *p*_d_ = 0.9, *p*_r_ = 0.1. **C** Slow degradation and fast recovery profile: *p*_d_ = 0.1, *p*_r_ = 0.9. **D** Slow degradation and slow recovery profile: *p*_d_ = 0.1, *p*_r_ = 0.1.

## Discussion

We have found an optimal time limit for the development of a new technology that anticipates the successes and failures of development and deployment. This R&D investment strategy is similar to an optimal stopping time, with a key difference: if the technology is ready before the end of this time limit, technology development investments stop. Determining this time limit allows decision-makers to reallocate their resources to other projects if the technology is not ready by then. Our analysis shows that this time limit depends on the belief in the future success of technology development, costs of development, and expected deployment benefits. By explicitly accounting for the structural uncertainty of R&D outcomes (or models for success or failure, see Methods), our approach fills a gap in the R&D literature, which typically assumes a known model structure for technology development. This concept, called model uncertainty [65; 20], can be incorporated into other methods for R&D resource allocation, such as Real Options Analysis [54], and accommodate other model structures for technology development [26; 64]. Our approach can significantly improve allocation of resources for R&D programs, in many domains, including biodiversity conservation, energy production and the technology industry more broadly.

We anticipate all possible future system responses to technology deployment using an AI approach (see Methods). This approach enables decision-makers to explicitly account for every possible system response to technology deployment, including beneficial and harmful consequences the technology might cause. Addressing uncertainties in technology deployment is essential to deploy new technologies as efficiently as possible, especially in domains where effects of technology remain uncertain despite experimentation, such as biodiversity conservation and geoengineering [47]. We demonstrate that these uncertain deployment outcomes are crucial for determining how long we can develop new technologies for (Fig. 3 and 4).

Our approach to finding the time limit for technology development is flexible and can be adapted to other technology deployment strategies. We assumed risk-neutrality when calculating the maximum expected benefits of technology deployment (*R*_AM_, see Methods), but our approach can easily accommodate risk-averse or risk-prone preferences [72; 4]. Adapting our approach to these alternatives would require first calculating annual expected benefits of deployment generated by the desired strategy (for example using robust optimisation [13] for a risk-averse strategy or a risk-seeking setting to obtain a risk-prone strategy [41]), and then changing the variable *R*_dep_ to these newly calculated expected benefits.

In an illustrative case study motivated by the technologies currently being developed for the Great Barrier Reef [50], we assumed that the development stage of the new technology and the health of an entire ecosystem can be categorized in a binary manner (idle or ready, and unhealthy or healthy respectively, see Methods and SI A and B). These are necessary simplifications of a technology development process that is continuous, with multiple development stages; and of an ecosystem that is spatially and ecologically complex. This simplification allowed us to derive critical insights into the key variables influencing the results (e.g., belief of future technology development success, decline or recovery probabilities, and uncertain effects of the technology). The strength of our modelling approach using POMDPs is the ability to readily expand the state and action spaces in future work and to leverage the extensive research in AI to solve POMDPs with large state and action spaces, such as Deep Reinforcement Learning [32; 21]. Increasing the number of states and actions increases model complexity but also enhances realism. For example, states space can be augmented to account for different levels of technology readiness and successive improvements, or for different ecological recovery phases and spatial heterogeneity of the ecosystem. However, greater model complexity leads to increased difficulty in interpreting and explaining results [30; 52], which can compromise trust and limit applications. While we strongly believe that the adaptive management framework is ideal for addressing uncertain development and deployment outcomes, alternative modelling approaches could have been chosen. For example, we opted for a model uncertainty formulation (POMDPs) rather than a parameter uncertainty formulation of the adaptive management problem (MDPs, [53; 34]). Historically, both formulations have presented strengths and weaknesses due to their computational complexity; however these considerations need to be reassessed in light of the emergence of deep reinforcement learning approaches [20; 17; 21].

We also assumed that the system dynamics are stationary, an assumption that becomes questionable if the dynamic system of interest operates in an environment that changes over the same timescale as the proposed technology development and deployment phases (e.g. climate change [71]). Growing pressures on ecosystems might push them past degradation tipping points, where restoration becomes too difficult or even impossible [69; 12; 24], or conversely, ecosystems might adapt and develop resistance to pressures [70]. Thus, our approach provides an upper bound – and conversely, a lower bound – of the time limit for technology development when we do not account for increased ecosystem degradation rates due to climate change (or, respectively, for an ecosystem adaptable to pressures). POMDP models can be adapted to account for non-stationary dynamics by setting transition probabilities between possible futures in a potential extension to this work [56; 57] (*T*_*y*_ in SI B.2).

In short, we have developed a computational decision-making approach that can determine how long to invest in technology development for dynamic systems. We have shown how the characteristics of a system can drastically change the technology development strategy, and thus inform optimal allocation of resources to maximise outcomes. Now more than ever, these approaches are needed to guide investments of limited resources for technology development and deployment in dynamic systems across a variety of disciplines [40; 66; 48]. In ecology, our approach guides strategic investments in new technologies to enhance the resilience and long-term health of iconic ecosystems like the Great Barrier Reef, advancing effective conservation efforts.

## Methods

We address the optimisation problem of when to stop investing in new technologies in two steps: technology development and technology deployment. To achieve this, we solve two adaptive management problems, both with the objective of achieving the maximum expected discounted sum of long-term benefits (ecosystem services) and costs (technology development and deployment). Adaptive management is a solution to manage systems when the outcomes of actions are uncertain [38; 73]. Adaptive management is particularly suitable for technology development and deployment since the future success of both steps is uncertain. In an adaptive management framework, a decision-maker first identifies possible alternative futures (or models) that will compete in the decision-making process. Optimal adaptive management strategies dynamically learn the most effective actions, given the level of confidence that each possible scenario represents the ground truth [29; 28; 65; 58]. We model each adaptive management problem using Partially Observable Markov Decision Processes (POMDPs, [7; 76]). POMDPs are mathematical models to optimise sequential decisions when the decision-maker can only partially observe the system. Formulating adaptive management problems as POMDPs guarantees optimality by finding the best trade-off between reducing uncertainties and gaining benefits using existing knowledge [19; 61; 20].

### Technology development

For clarity, we first introduce the simpler case where the probability of successful technology development is known, which we model as a Markov decision process (MDP). We then present the more realistic case considering possible models of technology development failure by including partial observability, with a POMDP model.

When we assume the technology development will succeed, we can determine the optimal investment strategy for technology development by balancing development costs and expected benefits of deployment. We model this decision problem using an infinite time-horizon 2-state MDP. Formally, let *X* = {idle, ready} represent the set of fully observable states describing the development progress of a new technology. We assume that the initial state is *x*_0_ = idle (i.e., the technology is not yet developed to a point that it is ready for deployment). The set of actions available is *A* = {surrender, invest in R&D (if technology is idle) or deploy (if technology is ready)}. Implementing the action *surrender* prevents the transition from *idle* to *ready* (*P* (ready|idle, surrender) = 0). Conversely, the action *invest in R&D* allows the technology to transition from *idle* to *ready* with probability *P* (ready|idle, invest in R&D). Here, we set *P* (ready|idle, invest in R&D) = 0.1 because the expected development time for large interventions for the case study inspiring our problem is 10 years [50]. Once *ready*, the technology cannot transition back to *idle*. Implementing an action in each state generates rewards and comes at a known cost, summarized in the reward function (*r*, see Table S2). When the action *surrender* is taken, the decision-maker receives a baseline reward (*R*_BAU_), representing the ecosystem services generated without the technology. This baseline reward depends on the degradation and recovery features of the managed ecosystem (see SI B.1). Taking the action *invest in R&D* in the state *idle* yields the same baseline rewards and incurs a development cost (*C*_dev_). Deploying the technology in the *ready* state provides rewards generated by the deployment of the new technology (*R*_dep_), that are expected, though not guaranteed, to exceed the baseline reward *R*_BAU_. We apply a discount factor (*γ <* 1) to weigh the relative importance of immediate rewards compared to future ones.

Solving an MDP means determining the best action to implement in each state to maximise the expected sum of discounted rewards over time [67]. While solving this MDP is straightforward, this formulation inadequately accounts for the possible failure of technology development as a solution may recommend investing in technology development indefinitely. To address this limitation, we improve upon this formulation by explicitly accounting for potential development failures by augmenting the state space with a set of unobservable possible future scenarios (or models) for technology development: *Y* = {success, failure}. These states describe an underlying characteristic of the technology - it is either viable and can successfully be developed, or it is unviable and will eventually fail to be developed. Incorporating these unobservable states for the technology converts the MDP into a POMDP, as the underlying dynamics are now unobservable. By including these future scenarios as a hidden state variable, we can now define the state transition probabilities with respect to each scenario (the rest of the model is unchanged). Under the future scenario *success*, the yearly probability of transitioning from *idle* to *ready* when *investing in R&D* is positive (*P* (ready|idle, invest in R&D, success) = *p*_dev_). Conversely, a failed technology development is a scenario where the yearly probability of transitioning from *idle* to *ready* when *investing in R&D* is null (*P* (ready|idle, invest in R&D, failure) = 0). Since the true development scenario is unknown to the decision maker, the development scenario needs to be inferred using past observations. In a POMDP, all the past decisions and observations are summarized into a sufficient statistic [15], called the belief state *b*, which is a probability distribution over *Y*. The initial belief state *b*_0_ represents the decision-maker’s initial confidence in the technology’s successful development. The belief state at time *t, b*_*t*_(success), represents the belief that the technology will be successfully developed in the future, while *b*_*t*_(failure) = 1 − *b*_*t*_(success) represents the belief of development failure.

The decision-making proceeds as follows. At each time step *t*, the state of the system is fully described by a technology development state (*x*_*t*_) and a belief state over the development scenarios (*b*_*t*_). The decision maker implements an action *a*_*t*_, and receives a reward that depends on the current state and the action chosen (*r*(*x*_*t*_, *a*_*t*_)). At the following time step *t*+1, the technology development state transitions from *x*_*t*_ to *x*_*t*+1_. The technology state itself is observable – the decision-maker can see whether the technology has been successfully developed (entered the *ready* state) or remains in the *idle* state. If the technology remains *idle*, the decision-maker uses that information to update their belief that development will ever be successful. The belief state is updated to *b*_*t*+1_ according to the resulting outcome *x*_*t*+1_ and implemented action *a*_*t*_ using Bayes’ rule:

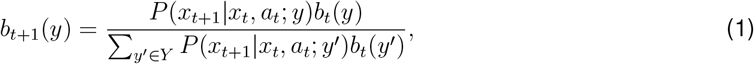

where *P* (*x*_*t*+1_|*x*_*t*_, *a*_*t*_; *y*) represents the probability of transitioning from a given state *x*_*t*_ to another state *x*_*t*+1_, when the action *a*_*t*_ is implemented, and assuming *y* is the true scenario.

Here, solving a POMDP means determining an optimal investment strategy to maximize the expected infinite discounted sum of rewards. An optimal strategy *π*^*∗*^ : *X × B → A* takes into account a combination of development stage of the technology (*idle* or *ready*) and belief over the possible development scenarios (*b*_*t*_) to give an investment decision recommendation (*surrender, invest in R&D* or *deploy*).

We discover that the optimal investment strategy has a well-defined structure. When the state *x* is *ready*, the optimal strategy is to strategically deploy the technology if *R*_dep_ *> R*_BAU_ (independently of the belief in successful technology development as when the technology is *ready* the outcomes of development investments are no longer a source of uncertainty). When the state *x* is *idle*, the decision to invest in R&D or surrender depends on the belief in successful technology development. Of interest to us to determine an analytical approximation of the time limit for technology development (*T*_max_) is the value of the belief in development success where the strategy switches from *invest in R&D* to *surrender* (*b*_*i/s*_). When the current belief in development success is greater than this switching belief threshold, the optimal action is to *invest in R&D*, and *surrender* when smaller. We approximate this switching belief threshold *b*_*i/s*_ building on [18] and [61] (see SI C for details):

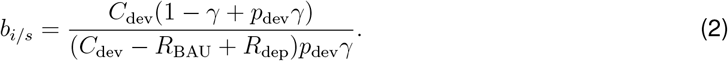

When the action *invest in R&D* is implemented and the technology does not transition from *idle* to *ready*, the belief in successful technology development decreases (see eq 1). From this expression of the switching belief threshold we derive the maximum number of years investing in technology development, *T*_max_, by finding the time it takes for the current belief in development *success* (*b*_*t*_(*y* = *success*), denoted *b*_*t*_ for simplicity) to decline to *b*_*i/s*_. The analytical approximation of *T*_max_ is then

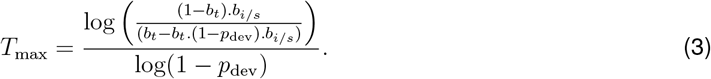

Note that this approximation can be used at any time step to determine the time left for technology development. This approximation gives the total time available for technology development (*T*_max_), when it is used with the initial belief in technology development success (*b*_0_, see Fig 2).

### Technology deployment

If a technology is successfully developed, the system’s response to technology deployment is still uncertain. We must develop an optimal plan for when to deploy the technology over many years. The outcome achieved by this optimisation gives the expected value of deploying that technology. We propose a novel approach to calculate the expected benefits of technology deployment (*R*_dep_) when technologies are designed for complex dynamic systems. Similar to the development stage, we plan technology deployment as an adaptive management problem, to account for the uncertain ecosystem’s response to technology deployment. Due to the complexity of defining this future response, we use the universal adaptive management solver, an AI algorithm that builds on mathematical properties of adaptive management problems, to determine a minimum but complete set of all possible future scenarios for technology deployment [60] (Table 1).

**Table 1:** Summary of all four possible outcomes of technology deployment identified by the universal adaptive management solver [60]. Each model represents the efficiency of the action *deploy* relative to the action *business as usual* for each state. See Fig. S4. for detailed parameterization of each model.

| Model | Optimal management action for each state |  |  |  |
| --- | --- | --- | --- | --- |
| | $m_1$ | $m_2$ | $m_3$ | $m_4$ |
| unhealthy | business as usual | business as usual | deploy | deploy |
| healthy | business as usual | deploy | business as usual | deploy |

We frame the technology deployment phase as an adaptive management problem using a POMDP (see SI B). Let *S* = {unhealthy, healthy} be the finite set of fully observable states, representing the possible health status of the ecosystem. We assume that a technology is ready for deployment, but the outcomes of deployment on each ecosystem state are unknown, while the effects of current business as usual management activities are assumed fully known. We therefore consider two actions, and ask which action is best to implement in each state (*U* = {business as usual, deploy}). We define the reward function so that it represents costs and benefits of implementing each action in each state (see Table S3). When *healthy*, the decision maker receives the ecosystem services generated by a healthy ecosystem (*B*_h_). Similarly, when *unhealthy*, the rewards received by the decision maker are the ecosystem services generated by an unhealthy ecosystem (*B*_u_). Implementing the action *business as usual* incurs no cost, while implementing the action *deploy* incurs a known deployment cost (*C*_deploy_). The dynamics of the ecosystem when implementing the action *business as usual* are assumed known (see SI B.1), and are characterized by *p*_r_ (the annual probability of recovering from unhealthy to healthy) and *p*_d_ (the annual probability of degrading from healthy to unhealthy). As the future effects of technology deployment are uncertain, we consider a non-observable set of possible outcomes of technology deployment (*M*), determined with the AI algorithm universal adaptive management solver [60]. Briefly, this algorithm derives the smallest discrete but complete set of all possible future scenarios that could occur. This algorithm outputs four possible outcomes for technology deployment, each representing the efficiency of the action *deploy* in each state relative to the action *business as usual* and *M* = {*m*_1_, *m*_2_, *m*_3_, *m*_4_} (see Table 1 and SI B.3). The true deployment scenario is unknown to decision makers, i.e, they cannot observe which model in *M* is the true model describing how the ecosystem will respond to the technology. As in the development model, the decision-makers here have a belief about which model is true (*β*), which updates each time an action is taken and the ecosystem response is observed (see eq. 1).

The decision-making is similar to the technology development POMDP, with an ecosystem state (*s*_*t*_) and a belief over possible ecosystem responses to technology deployment (*β*_*t*_). The optimal strategy maximises the expected sum of long-term benefits (value function *W* ^*∗*^).

For analysis purposes, we summarize the belief state *β*_*t*_ with two marginal belief states 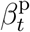 and 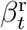. 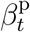 represents the belief that technology deployment is the optimal action if the ecosystem is healthy (for preventing degradation from healthy to unhealthy). 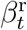 represents the belief that technology deployment is the optimal action if the ecosystem is unhealthy (for restoration from unhealthy to healthy).

By solving this adaptive management problem, we optimally calculate the expected benefits of technology deployment for managing complex dynamic systems when the deployment effects are uncertain. We set *R*_dep_ as the yearly benefits of technology deployment with adaptive management *R*_AM_, which we obtain by annualizing the expected long-term benefits of technology deployment (*R*_AM_ = (1−*γ*)*W* ^*∗*^ see SI B.2). Code and results are available on Git-hub: https://github.com/luzvpascal/technoDev.

## Acknowledgements

We thank Dr. Ken Anthony for providing insightful feedback on this study. LVP and KJH were funded by Australian Research Council Discovery Early Career Researcher Award DE200101791. MPA’s contribution was funded by an Australian Research Council Discovery Early Career Resarcher Award DE200100683. LVP, KJH and MPA acknowledge funding from the ARC SRIEAS Grant SR200100005 Securing Antarctica’s Environmental Future.

