## Supplemental material for "Developing new technologies to protect ecosystems: planning with adaptive management"

#### A Model: POMDP for technology development

We formally define a POMDP for planning technology development as a tupple $\langle X, Y, b_0, A, T, O, Z, r, H, \gamma \rangle$ with:

- a finite set of fully observable states $X$. $X$ is the combination of the possible development states of the project candidate $X = \{$idle, ready$\}$;
- a finite set of actions $A$, which contains all the possible decisions available at each time step. $A = \{$surrender, investing in R&D (if technology idle), deploying the technology (if technology ready)$\}$;
- a finite set of non-observable states $Y$. In our model, $Y$ represents the possible scenarios for technology development outcomes: $Y = \{$success, failure$\}$. When undertaking R&D, a major concern lies in the uncertainty surrounding the outcomes of R&D processes and whether and when

a development investment will yield a deployable new technology. Under the future scenario *success*, the yearly probability of transitioning from *idle* to *ready* when *investing in R&D* is positive ($P(\text{ready}|\text{idle}, \text{invest in R\&D}, \text{success}) = p_{\text{dev}}$). Conversely, a failed technology development is a scenario where the yearly probability of transitioning from *idle* to *ready* when *investing in R&D* is null ($P(\text{ready}|\text{idle}, \text{invest in R\&D}, \text{failure}) = 0$). We detail the parameters of the possible scenarios for project development, see Figure S1. Because the elements of $Y$ are not observable or measurable, they need to be inferred using past observations. We represent this past information as a probability distribution over all the elements of $Y$, which is called a belief state $b$. The initial belief state is $b_0$;

- $T : X \times Y \times A \times X \times Y \rightarrow [0, 1]$ is the state transition function, and represents the stochastic dynamics of the technology development. We define $T(x, y, a, x', y') = P(x'|x, y, a)P(y'|y)$. Here, $P(x'|x, y, a)$ is the probability that the technology transitions from $x$ to $x'$ when the action $a$ is implemented and according to the model $y$, which is a Markov chain (see Figure S1). $P(y'|y)$ represents the transition probability between the elements of $Y$, and gives the probability that the development scenario transitions from $y$ to $y'$. This function relates to the overall stationarity of the system. In this work, we assume that the system is stationary, and $P(y'|y) = 1$ if $y' = y$ and 0 otherwise.

- The states of the system $X$ are fully observable, which means that decision makers know the development stage of the technology at every time step and without uncertainty. Then, the set of observations $O$ is the same as the set of states $X$. The observation function $Z$ gives the probability we obtain an observation $o$ given a state $x$. Thus, here $Z$ is the identity function.

- The reward function $r : X \times A \rightarrow \mathbb{R}$ returns the benefits and costs of executing an action $a$ in a given state. We propose here a unitless reward function normalized according to the maximum possible benefits. We summarize variables defining the reward function $r$ in Table S1. If the action *surrender* is implemented, the decision-maker perceives a baseline reward corresponding to business-as-usual interventions ($R_{\text{BAU}}$, see Section B.1), when the technology is *idle* or *ready*. If the decision-maker decides to invest in R&D (only possible if the technology is *idle*), then they perceived the baseline reward deduced by the costs of technology development ($R_{\text{BAU}} - C_{\text{dev}}$). If the technology is *ready* and the decision-maker decides to *deploy*, then we the decision-maker perceives the rewards achieved by an deployment strategy ($R_{\text{dep}}$, that can be set to $R_{\text{AM}}$ as in Section B.2). The link between technology development and technology deployment appears in the reward function through $R_{\text{dep}}$. A technology successfully developed (a ready technology) is not necessarily efficient for management, and its deployment on-ground also needs to be planned (see Section B.2).

- $H$ represents the time horizon of the problem. Here we assume $H = +\infty$ as we aim to relate the long term costs and benefits of investment strategies.

- $\gamma \in [0, 1]$ is the discount factor. $\gamma$ relates the value of future rewards compared to their present value: a discount factor lower than one indicates that rewards are more valuable in the present than in the future. We set $\gamma = 0.9$.

Here, a strategy $\pi : X \times B \to A$ is a function mapping a state $(x, b)$ to an action $a = \pi^*(x, b)$. An optimal strategy $\pi^*$ is a strategy that maximises the criterion:

$$V^*(x_0, b_0) = \max_{\pi} \mathbb{E}\left(\sum_{t=0}^{+\infty} \gamma^t R(x_t, b_t, \pi(x_t, b_t)) \middle| x_0, b_0\right) = \mathbb{E}\left(\sum_{t=0}^{+\infty} \gamma^t R(x_t, b_t, \pi^*(x_t, b_t)) \middle| x_0, b_0\right), \quad \text{(S1)}$$

where $R : X \times B \times A \to \Re$ is the reward function over states and beliefs. Essentially, $R(x, b, a)$ is the immediate reward received when the current state is $(x, b)$ and the chosen action is $a$. Formally $R(x, b, a) = \sum_{y \in Y} r(x, a)b(y) = r(x, a)$. The function $r : X \times A \to \Re$ is the immediate reward function described by Table S2.

### B Ecosystem services under current management and with adaptive technology deployment

#### B.1 Ecosystem services under current management and current ecosystem dynamics

We model the Great Barrier Reef as a stochastic and dynamic system, that can transition between a healthy or an unhealthy state. As the Australian Institute of Marine Science, we use hard coral cover as an indicator of reef health [2; 6; 7], and define a healthy state as above 30% hard coral cover [7]. We estimate the transition function of the reef ecosystem under current business as usual activities (a Markov chain) using existing data collected and analysed by the Australian Institute of Marine Science [3], and publicly available on the ReefCloud platform [1]. ReefCloud is an online platform supported by the Australian Institute of Marine Science and the Australian Government, and it uses machine learning and statistical models to estimate coral cover at different spatial scales based on long term monitoring reef images across the Great Barrier Reef. To estimate the transition function of the reef ecosystem, we use the hard coral cover predictions between 2004 and 2022 produced by ReefCloud, for the Central and Southern Great Barrier Reef (see Figure S2**A**). We categorize the coral cover value as healthy (above 30%) or unhealthy (under 30 %) (see Figure S2**B**), and we estimate the corresponding Markov chain using the R-package `markovchain` [11]. We obtain the Markov chain described in Figure S2**C**. We estimate the annual transition probabilities of degradation from healthy to unhealthy reef at 0.8 per year ($P(\text{unhealthy}|\text{healthy, BAU}) = 0.8$), and recovery from unhealthy to healthy at 0.2 per year ($P(\text{healthy}|\text{unhealthy, BAU}) = 0.2$). Because this time series is short, the estimation of the Markov chain parameters are highly uncertain and we explore how variations of these two parameters influence the general technology development and deployment results (see Fig S7).

We compute the expected benefits of current management ($R_{\text{BAU}}$) as follows. When the Great Barrier Reef is healthy, it generates a range of ecosystem services worth $B_{\text{h}}$ =$6.4 billion AUD ($4.3 billion USD) yearly [8]. We estimate the ecosystem services produced in an unhealthy state by assuming that ecosystem services are proportional to the coral cover. According to [8], the Great Barrier Reef generated 6.4 billion AUD ($4.3 billion USD) in the financial year 2015-2016. In 2015, the coral cover was estimated at 32% (corresponding to a *healthy* state). We estimate the average coral cover when the Great Barrier Reef is categorized as *unhealthy* to be 21% (between 2004 and 2022, see Fig S2). Proportionally, we estimate that the ecosystem services produced when the ecosystem is unhealthy ($B_{\text{u}}$) are 67% of the healthy value. At each time step, the expected produced ecosystem services, $R_{\text{BAU}}$, are the sum of ecosystem services produced in each state weighted by the state-probabilities of the Markov chain's steady state. For example, for our case study, the steady state for the Markov chain is $[P(\text{healthy}) = 0.2; P(\text{unhealthy}) = 0.8]$, therefore the expected ecosystem services under current management are $R_{\text{BAU}} = 0.2B_{\text{h}} + 0.8B_{\text{u}}$.

#### B.2 Model: POMDP for technology deployment

In this section, we detail how we calculate the expected benefits of new technology deployment with adaptive management, $R_{\text{AM}}$, under the hypothesis that the new technology is successfully developed (we set $R_{\text{dep}} = R_{\text{AM}}$ in the POMDP for technology development). To estimate $R_{\text{AM}}$, we undertake an adaptive management approach under model uncertainty, and find an optimal management strategy by posing and solving this adaptive management problem as a POMDP [5]. Because we aim to account for all possible future scenarios for technology deployment, we use the 2-state n-action universal adaptive management solver presented in [9]. The universal adaptive management solver determines a small set of possible dynamics scenarios with the warranty to include all possible futures for technology deployment.

We determine $R_{\mathrm{AM}}$ by posing the deployment of new technologies for ecosystem management as a POMDP.

- The set of fully observable states is $S = \{\text{unhealthy, healthy}\}$, describing the possible states of the ecosystem (as in Section B.1).

- The set of possible outcomes for technology deployment (non-observable) is $M$. We determine $M$ using the universal adaptive management solver (see Section B.3) and obtain 4 possible scenarios :

$$M = \left\{\begin{array}{l} m_1\text{: BAU is the optimal action if the ecosystem is healthy or unhealthy} \\ m_2\text{: Technology deployment is the optimal action if the ecosystem is healthy only, BAU if unhealthy} \\ m_3\text{: Technology deployment is the optimal action if the ecosystem is unhealthy only, BAU if unhealthy} \\ m_4\text{: Technology deployment is the optimal action if the ecosystem is healthy or unhealthy} \end{array}\right\}.$$

  We summarize the obtained scenarios for the Great Barrier Reef case study in Figure S4. Because the elements of $M$ are not observable, we infer them at each time step using a sufficient statistic called the belief state $\beta$. A belief state is a probability distribution among the elements of $M$, and $\beta_t(m)$ is the probability that the true scenario is $m$ at time step $t$. We summarize the belief state $\beta_t$ with two marginal beliefs states $\beta_t^{\mathrm{p}}$ and $\beta_t^{\mathrm{r}}$. $\beta_t^{\mathrm{p}}$ represents the belief that technology deployment is the optimal action if the ecosystem is healthy (for preventing degradation from healthy to unhealthy). $\beta_t^{\mathrm{r}}$ represents the belief that technology deployment is the optimal action if the ecosystem is unhealthy (for restoration from unhealthy to healthy). Formally:

$$\beta_t^{\mathrm{p}} = \beta_t(m_2) + \beta_t(m_4),$$

$$\beta_t^{\mathrm{r}} = \beta_t(m_3) + \beta_t(m_4);$$

- The set of actions is $U = \{$business as usual, deploy$\}$;

- The reward $r_{\mathrm{reef}}$ function is defined Table S3. When the ecosystem is *healthy* (resp. *unhealthy*), the decision-maker perceives a reward corresponding to the ecosystem services generated by a healthy (resp. unhealthy) ecosystem ($B_{\mathrm{h}}$ and $B_{\mathrm{u}}$ respectively). The action *business as usual* incurs no cost, while the action *deploy* incurs a technology deployment cost ($C_{\mathrm{deploy}}$);

- The stochastic dynamics of the system are $T_{\mathrm{reef}} : S \times U \times S \times M \rightarrow [0, 1]$. For example $T_{\mathrm{reef}}($unhealthy, deploy, healthy$, m_1)$ represents the probability that the reef transitions from an unhealthy to a healthy state when the action "deploy" is implemented, and according to the scenario $m_1$. Here we assume that the dynamics of the system for the action *business as usual* are fully known by the decision-maker (see Section B.1), and are the same across all models in $M$;

- Because we assume only the current health state of the reef is observable, the set of observations is $S$, and the observation function is the identity matrix.

- $H$ represents the time horizon the of problem. Here we assume $H = +\infty$ as we aim to relate the long term costs and benefits of strategies.

- $\gamma \in [0, 1]$ is the discount factor. $\gamma$ relates the value of future rewards compared to their present value: a discount factor lower than one indicates that rewards are more valuable in the present than in the future. We set $\gamma = 0.9$.

The value function of this POMDP is $W^*$, which represents the maximum expected sum of discounted rewards obtained when deploying a new technology with an adaptive management program. Here, $\delta : S \times \mathbb{B} \rightarrow U$ is a strategy mapping a state and a belief state to an action.

$$W^*(s, \beta) = \max_{\delta} \mathbb{E} \left( \sum_{t=0}^{+\infty} \gamma^t r_{\mathrm{reef}}(s_t, \delta(s_t, \beta_t)) | s_0 = s, \beta_0 = \beta \right).$$

$R_{\mathrm{AM}}$ is the yearly expected benefits of undertaking an adaptive management approach for technology deployment. We compute $R_{\mathrm{AM}}$ as:

$$R_{\mathrm{AM}} = [W^*(\text{healthy}, \beta_0) * P(\text{healthy}) + W^*(\text{unhealthy}, \beta_0) * P(\text{unhealthy})]\,(1-\gamma),$$

where $P(\text{healthy})$ and $P(\text{unhealhy})$ are the steady states of the Markov chain describing the reef dynamics under current management (Section B.1). Multiplying by $(1-\gamma)$ allows to transform the long-term sum of discounted rewards into yearly expected rewards.

##### B.3 Universal adaptive management solver and technology deployment scenarios

The universal adaptive management solver [9] operates in four steps, as illustrated in Figure S3. First, the user inputs a reward function, $r$. The algorithm identifies the infinite set of possible candidate scenarios for that reward function, which are all the possible transition functions of a Markov Decision Process with a reward function $r$. Second, the algorithm samples scenarios using a uniform distribution, and classifies them according to their optimal policy. Finally, the algorithm evaluates the average scenarios of each possible optimal policy: these are referred to as **universal models**, and are the outputs of the algorithm. In this study, we adapt this algorithm to incorporate our current knowledge about the system dynamics when the action *business as usual* is implemented (Section B.1).

#### C Analytical approximation of maximum number of years investing in project development

In this section, we detail how we approximate analytically the optimal time limit for technology development. Our approach builds on an analytical approximation of $V^*$ the value function of the technology development POMDP (Section C.1). This allows to find an approximation of $b_{i/s}$, the belief in project feasibility where the optimal policy changes from invest to surrender (Section C.2). The analytical expression of $b_{i/s}$ allows to determine analytically the maximum number of years investing in project development, when starting from any belief in future technology development success ($b_t(\text{success})$), denoted $b_t$ for simplicity.

#### C.1 Value function approximation of technology development POMDP

We build on [10], to provide an analytical lower bound approximation of $V^*$, the value function of the POMDP for technology development (Figure S5). To build this analytical lower bound, we derive $V_i$ (resp. $V_s$), the sum of expected rewards when the policy *invest in R&D* (resp. *surrender*) is implemented. Following the procedure described in [10], we evaluate $V_i$ and $V_s$ in the corners of the belief space $B$ i.e. when the decision maker knows if technology development will be successful or not. Peron et al. [10] demonstrate that the interpolation between $V_i(\text{idle}, b = 1)$ (the expected benefits of investing in R&D when the technology is *idle*, and the technology development will be successful) and $V_i(\text{idle}, b = 0)$ (the expected benefits of investing in R&D when the technology is *idle*, and the technology development will fail) is a lower bound of $V^*(\text{idle}, b)$ for any $b$ (red line in Figure S5) – Likewise between $V_s(\text{idle}, b = 1)$ and $V_s(\text{idle}, b = 0)$ (purple line in Figure S5). Peron et al. [10] show that the point wise maximum of these two interpolations is a lower bound of $V^*$ (blue line in Figure S5), and moreover, $V^*(\text{idle}, b = 1) = V_i(\text{idle}, b = 1)$ and $V^*(\text{idle}, b = 0) = V_s(\text{idle}, b = 0)$. We approximate the belief in future successful technology development where the optimal strategy switches from *invest in R&D* to *surrender* ($b_{i/s}$) as the intersection of these two interpolations (intersection of red and purple lines in Figure S5).

We derive $V_s$ and $V_i$ analytically to determine $b_{i/s}$ analytically, and then the time limit for technology development.

The value of implementing the strategy *surrender* $V_s$ depends on the rewards obtained when implementing the action *surrender* ($R_{\text{BAU}}$) and $V_s$ does not depend on the belief in successful technology development.

$$V_s(\text{idle}, b = 1) = V_s(\text{idle}, b = 0) = \sum_{t\geq 0}(\gamma^t R_{\text{BAU}}) = \frac{R_{\text{BAU}}}{1-\gamma}. \tag{S2}$$

Let us derive the expected sum of discounted rewards when implementing the strategy *invest in R&D* knowing that technology development will fail: $V_i(\text{idle}, b = 0)$. We recall that $C_{\text{dev}}$ is the relative cost of investing in R&D per year. Then by continuously investing in technology development although technology development will never be successful, we obtain

$$V_i(\text{idle}, b = 0) = \sum_{t\geq 0}(\gamma^t (R_{\text{BAU}} - C_{\text{dev}})) = \frac{R_{\text{BAU}} - C_{\text{dev}}}{1-\gamma}. \tag{S3}$$

Similarly, if the technology is successfully developed, the expected sum of discounted benefits from adaptive technology deployment are:

$$V_i(\text{ready}, b=1) = R_{\text{AM}} \sum_{i=0}^{\infty} \gamma^i = \frac{R_{\text{AM}}}{1-\gamma}. \quad \text{(S4)}$$

We derive the expected sum of discounted rewards when implementing the strategy *invest in R&D* knowing that technology development will succeed ($V_i(\text{idle}, b=1)$) by backward dynamic programming, as summarized in Table S4. To do so, we analytically derive the value function of the technology development POMDP until time $t$:

$$V_i^t(\text{idle}, b=1) = \mathbb{E}\left(\sum_{k=0}^{t} \gamma^k r(x_k, \pi(x_k, b_k)) | x_0 = x, b=1\right)$$
$$\text{s.t. } \pi(\text{idle}, b) = \text{invest in R\&D}, \quad \pi(\text{ready}, b) = \text{deploy}, \forall b$$

We demonstrate by induction that at time $t \geq 1$, the value function $V_i^t$ is:

$$V_i^t(\text{idle}, b=1) = (R_{\text{BAU}} - C_{\text{dev}}) \left(\sum_{i=0}^{t} (1-p_{\text{dev}})^i \gamma^i\right) + (1-p_{\text{dev}})^t \gamma^t R_{\text{BAU}} \quad \text{(S5)}$$
$$+ p_{\text{dev}} \gamma R_{\text{AM}} \left(\sum_{i=0}^{t-1} (1-p_{\text{dev}})^i \gamma^i [\sum_{j=0}^{t-i-1} \gamma^j]\right). \quad \text{(S6)}$$

**Proof:**

First, for $t = 0$:

$$V_i^0(\text{idle}, b=1) = R_{\text{BAU}}, \quad V_i^0(\text{ready}, b=1) = R_{\text{AM}}.$$

For $t = 1$, using Bellman's equation of Dynamic Programming [4]:

$$V_i^1(\text{idle}, b=1) = \underbrace{R_{\text{BAU}} - C_{\text{dev}}}_{\text{immediate rewards}} + \underbrace{(1-p_{\text{dev}})\gamma R_{\text{BAU}}}_{\text{expected rewards if development fails}} + \underbrace{p_{\text{dev}}\gamma R_{\text{AM}}}_{\text{expected rewards if development succeeds}}$$

We now prove it for any $t$. We assume Eq. S5 is true for some $t \in \mathbb{N}$. Let us show that Eq. S5 is true for $t+1$.

$$V_i^{t+1}(\text{idle}, b=1) = R_{\text{BAU}} - C_{\text{dev}} + \underbrace{\gamma(1-p_{\text{dev}})V_i^t(\text{idle}, b=1)}_{\text{expected rewards if development fails}} + \underbrace{\gamma p_{\text{dev}} V_i^t(\text{ready}, b=1)}_{\text{expected rewards if development succeeds}}$$

We replace $V_i^t(\text{idle}, b=1)$ and $V_i^t(\text{ready}, b=1)$ using Eq. S5 and S4

$$\begin{aligned} V_i^{t+1}(\text{idle}, b=1) = {} & R_{\text{BAU}} - C_{\text{dev}} \\ & + \gamma(1-p_{\text{dev}})\Bigg[(R_{\text{BAU}} - C_{\text{dev}})\left(\sum_{i=0}^{t}(1-p_{\text{dev}})^i\gamma^i\right) \\ & + (1-p_{\text{dev}})^t\gamma^t R_{\text{BAU}} + p_{\text{dev}}\gamma R_{\text{AM}}\left(\sum_{i=0}^{t-1}(1-p_{\text{dev}})^i\gamma^i[\sum_{j=0}^{t-i-1}\gamma^j]\right)\Bigg] \\ & + \gamma p_{\text{dev}}\sum_{i=0}^{t} R_{\text{AM}}\gamma^i \end{aligned}$$

We rearrange this equation to show that Eq. S5 is true at $t+1$

$$\begin{aligned} V_i^{t+1}(\text{idle}, b=1) = {} & (R_{\text{BAU}} - C_{\text{dev}})\gamma^0(1-p_{\text{dev}})^0 \\ & + (R_{\text{BAU}} - C_{\text{dev}})\left(\sum_{i=1}^{t+1}(1-p_{\text{dev}})^i\gamma^i\right) \\ & + (1-p_{\text{dev}})^{t+1}\gamma^{t+1}R_{\text{BAU}} \\ & + p_{\text{dev}}\gamma R_{\text{AM}}\left(\sum_{i=1}^{t}(1-p_{\text{dev}})^i\gamma^i[\sum_{j=0}^{t-i-1}\gamma^j]\right) \\ & + \gamma p_{\text{dev}} R_{\text{AM}}(1-p_{\text{dev}})^0\gamma^0\sum_{i=0}^{t}\gamma^i \\ V_i^{t+1}(\text{idle}, b=1) = {} & (R_{\text{BAU}} - C_{\text{dev}})\left(\sum_{i=0}^{t+1}(1-p_{\text{dev}})^i\gamma^i\right) \\ & + (1-p_{\text{dev}})^{t+1}\gamma^{t+1}R_{\text{BAU}} \\ & + p_{\text{dev}}\gamma R_{\text{AM}}\left(\sum_{i=0}^{t}(1-p_{\text{dev}})^i\gamma^i[\sum_{j=0}^{t-i}\gamma^j]\right), \end{aligned}$$

which demonstrates that at time $t \geq 1$, the value function $V_i^t$ is:

$$V_i^t(\text{idle}, b=1) = (R_{\text{BAU}} - C_{\text{dev}}) \left( \sum_{i=0}^{t} (1-p_{\text{dev}})^i \gamma^i \right) + (1-p_{\text{dev}})^t \gamma^t R_{\text{BAU}} + p_{\text{dev}} \gamma R_{\text{AM}} \left( \sum_{i=0}^{t-1} (1-p_{\text{dev}})^i \gamma^i [\sum_{j=0}^{t-i-1} \gamma^j] \right).$$

**End of proof.**

To evaluate $V_i(\text{idle}, b=1)$ (expected sum of discounted benefits over an infinite horizon), we evaluate Eq. S5 when $t$ tends to $+\infty$. We first rearrange Eq. S5 to obtain an expression of $V_i(\text{idle}, b=1)$ that can be easily evaluated when $t$ tends to $+\infty$:

$$V_i^t(\text{idle}, b=1) = (R_{\text{BAU}} - C_{\text{dev}}) \left( \sum_{i=0}^{t} (1-p_{\text{dev}})^i \gamma^i \right) + (1-p_{\text{dev}})^t \gamma^t R_{\text{BAU}} + p_{\text{dev}} \gamma R_{\text{AM}} \left( \sum_{i=0}^{t-1} (1-p_{\text{dev}})^i \gamma^i [\sum_{j=0}^{t-i-1} \gamma^j] \right)$$

we simplify the sum expressions

$$V_i^t(\text{idle}, b=1) = (R_{\text{BAU}} - C_{\text{dev}}) \frac{1-(1-p_{\text{dev}})^{t+1}\gamma^{t+1}}{1-(1-p_{\text{dev}})\gamma} + (1-p_{\text{dev}})^t \gamma^t R_{\text{BAU}} + p_{\text{dev}} \gamma R_{\text{AM}} \left( \sum_{i=0}^{t-1} (1-p_{\text{dev}})^i \gamma^i \frac{1-\gamma^{t-i}}{1-\gamma} \right)$$

we distribute $(1-p_{\text{dev}})^i \gamma^i$

$$V_i^t(\text{idle}, b=1) = (R_{\mathrm{BAU}} - C_{\mathrm{dev}})\frac{1-(1-p_{\mathrm{dev}})^{t+1}\gamma^{t+1}}{1-(1-p_{\mathrm{dev}})\gamma}$$
$$+ (1-p_{\mathrm{dev}})^t\gamma^t R_{\mathrm{BAU}}$$
$$+ p_{\mathrm{dev}}\gamma R_{\mathrm{AM}}\left(\sum_{i=0}^{t-1}\frac{(1-p_{\mathrm{dev}})^i\gamma^i}{1-\gamma} - \frac{(1-p_{\mathrm{dev}})^i\gamma^t}{1-\gamma}\right)$$

we simplify the sum expression

$$V_i^t(\text{idle}, b=1) = (R_{\mathrm{BAU}} - C_{\mathrm{dev}})\frac{1-(1-p_{\mathrm{dev}})^{t+1}\gamma^{t+1}}{1-(1-p_{\mathrm{dev}})\gamma} + (1-p_{\mathrm{dev}})^t\gamma^t R_{\mathrm{BAU}}$$
$$+ p_{\mathrm{dev}}\gamma R_{\mathrm{AM}}\left(\frac{1-(1-p_{\mathrm{dev}})^t\gamma^t}{(1-\gamma)(1-(1-p_{\mathrm{dev}})\gamma)} - \gamma^t\frac{1-(1-p_{\mathrm{dev}})^t}{(1-\gamma)(1-(1-p_{\mathrm{dev}}))}\right)$$

We evaluate the limit of $V_i^t(\text{idle}, b=1)$ as $t \to +\infty$:

$$V_i(\text{idle}, b=1) = \lim_{t\to+\infty} V_i^t(\text{idle}, b=1)$$
$$V_i^t(\text{idle}, b=1) = (R_{\mathrm{BAU}} - C_{\mathrm{dev}})\frac{1-(1-p_{\mathrm{dev}})^{t+1}\gamma^{t+1}}{1-(1-p_{\mathrm{dev}})\gamma} + (1-p_{\mathrm{dev}})^t\gamma^t R_{\mathrm{BAU}}$$
$$+ p_{\mathrm{dev}}\gamma R_{\mathrm{AM}}\left(\frac{1-(1-p_{\mathrm{dev}})^t\gamma^t}{(1-\gamma)(1-(1-p_{\mathrm{dev}})\gamma)} - \gamma^t\frac{1-(1-p_{\mathrm{dev}})^t}{(1-\gamma)(1-(1-p_{\mathrm{dev}}))}\right)$$

these highlighted terms tend to 0 as $t \to +\infty$

$$V_i(\text{idle}, b=1) = \frac{R_{\mathrm{BAU}} - C_{\mathrm{dev}}}{1-(1-p_{\mathrm{dev}})\gamma} + \frac{p_{\mathrm{dev}}\gamma R_{\mathrm{AM}}}{(1-(1-p_{\mathrm{dev}})\gamma)(1-\gamma)}$$
$$V_i(\text{idle}, b=1) = \frac{R_{\mathrm{BAU}} - C_{\mathrm{dev}} + p_{\mathrm{dev}}\gamma R_{\mathrm{AM}}/(1-\gamma)}{1-(1-p_{\mathrm{dev}})\gamma}.$$

Therefore, the expected sum of discounted rewards obtained when applying the strategy *invest in R&D* is :

$$V_i(\text{idle}, b=1) = \frac{R_{\mathrm{BAU}} - C_{\mathrm{dev}} + p_{\mathrm{dev}}\gamma R_{\mathrm{AM}}/(1-\gamma)}{1-(1-p_{\mathrm{dev}})\gamma}. \quad \text{(S7)}$$

#### C.2 Approximation of belief state

Here we detail how we approximate the belief state $b_{i/s}$, where the optimal strategy switches from investing in project development to surrender.

We solve for $b_{i/s}$, $V_i(\text{idle}, b=1) \cdot b_{i/s} + V_i(\text{idle}, b=0) \cdot (1-b_{i/s}) = V_s(\text{idle}, b=1)$.

$$V_i(\text{idle}, b=1) \cdot b_{i/s} + V_i(\text{idle}, b=0) \cdot (1-b_{i/s}) = V_s(\text{idle}, b=1)$$

using Eq. S2, S3 and S7, we obtain

$$\Leftrightarrow \frac{R_{\text{BAU}} - C_{\text{dev}} + p_{\text{dev}}\gamma R_{\text{AM}}/(1-\gamma)}{1-(1-p_{\text{dev}})\gamma} b_{i/s} + \frac{R_{\text{BAU}} - C_{\text{dev}}}{1-\gamma}(1-b_{i/s}) = \frac{R_{\text{BAU}}}{1-\gamma}$$

$$\Leftrightarrow b_{i/s} = \frac{C_{\text{dev}}(1-\gamma+p_{\text{dev}}\gamma)}{(C_{\text{dev}} - R_{\text{BAU}})p_{\text{dev}}\gamma + p_{\text{dev}}\gamma R_{\text{AM}}}.$$

We approximate the belief state $b_{i/s}$, where the optimal strategy switches from investing in project development to surrender as:

$$b_{i/s} = \frac{C_{\text{dev}}(1-\gamma+p_{\text{dev}}\gamma)}{(C_{\text{dev}} - R_{\text{BAU}} + R_{\text{AM}})p_{\text{dev}}\gamma}.$$

#### C.3 Belief in project viability after n years investing in R&D

To determine the number of years necessary to keep investing in R&D before surrendering, we find a general formula for the belief state in successful technology development after $n$ years investing in technology development. We show here by induction that after investing for $n$ years in R&D, while not obtaining a ready technology, a belief state $b$ becomes:

$$b(\text{success})^{\text{n investments, idle}} = b_n = \frac{b \cdot (1-p_{\text{dev}})^n}{b \cdot (1-p_{\text{dev}})^n + 1 - b}. \tag{S8}$$

**Proof:** We demonstrate this assertion for one time investment $n = 1$:

$$b_1 = \frac{P(\text{idle}|\text{idle, invest in R\&D;success})b_0}{P(\text{idle}|\text{idle, invest in R\&D;success})b_0 + P(\text{idle}|\text{idle, invest in R\&D;failure})(1-b_0)}$$

$$b_1 = \frac{(1-p_{\text{dev}}).b_0}{(1-p_{\text{dev}})b_0 + 1(1-b_0)}.$$

We now prove it for any $n$. We assume Eq. S8 is true for some $n \in \mathbb{N}$. Let us show that Eq. S8 is true for $n+1$. We set $(1-p_{\text{dev}}) = p$:

$$\begin{aligned}
b_{n+1} &= \frac{P(\text{idle}|\text{idle, invest in R\&D;success})b_n}{P(\text{idle}|\text{idle, invest in R\&D;success})b_n + P(\text{idle}|\text{idle, invest in R\&D;failure})(1-b_n)} \\
&= \frac{p.b_n}{p.b_n + 1.(1-b_n)} \\
&= \frac{p.\frac{b.p^n}{b.p^n+1-b}}{p.\frac{b.p^n}{b.p^n+1-b} + 1 - \frac{b.p^n}{b.p^n+1-b}} \\
&= \frac{p.b.p^n}{p.b.p^n + b.p^n + 1 - b - b.p^n} \\
&= \frac{b.p^{n+1}}{b.p^{n+1} + 1 - b} \\
b_{n+1} &= \frac{b.(1-p_{\text{dev}})^{n+1}}{b.(1-p_{\text{dev}})^{n+1} + 1 - b}
\end{aligned}$$

**End of proof.**

#### C.4 Maximum number of years investing in project development

To find the maximum number of years investing in R&D, we solve for $n$, $b_n \geq b_{i/s}$. For simplicity, we keep our solution using $b_{i/s}$ instead of its analytical expression.

$$
\begin{aligned}
& b_n \geq b_{i/s} \\
& \frac{b.(1-p_{\text{dev}})^n}{b.(1-p_{\text{dev}})^n + 1 - b} \geq b_{i/s} \\
& b.(1-p_{\text{dev}})^n \geq (b.(1-p_{\text{dev}})^n + 1 - b) b_{i/s} \\
& (1-p_{\text{dev}})^n (b - b.(1-p_{\text{dev}}).b_{i/s}) \geq (1-b).b_{i/s} \\
& (1-p_{\text{dev}})^n \geq \frac{(1-b).b_{i/s}}{(b - b.(1-p_{\text{dev}}).b_{i/s})} \\
& n \log(1-p_{\text{dev}}) \geq \log\left(\frac{(1-b).b_{i/s}}{(b - b.(1-p_{\text{dev}}).b_{i/s})}\right) \\
& n \leq \frac{\log\left(\frac{(1-b).b_{i/s}}{(b - b.(1-p_{\text{dev}}).b_{i/s})}\right)}{\log(1-p_{\text{dev}})}
\end{aligned}
$$

Therefore, from any belief state $b$, we approximate the time limit for technology development investments as:

$$
T_{\max} = \frac{\log\left(\frac{(1-b).b_{i/s}}{(b - b.(1-p_{\text{dev}}).b_{i/s})}\right)}{\log(1-p_{\text{dev}})}.
$$

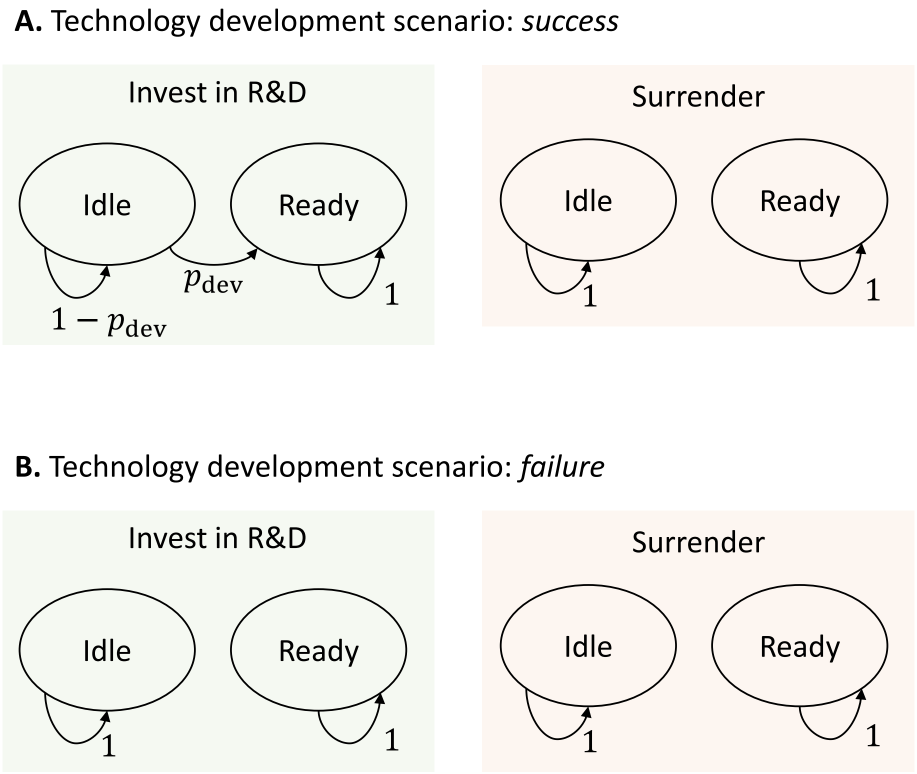


Figure S1: Dynamics of technology development for two possible scenarios (success and failure). For each scenario, we represent on the left-hand side the Markov chain describing the dynamics of technology development when the action *invest in R&D* is implemented, and on the right-hand side when the action *surrender* is implemented. In both scenarios, the technology cannot transition from *idle* to *ready* when the action *surrender* is implemented. (A) Scenario for successful technology development: the probability that the technology transitions from *idle* to *ready* when investing in R&D ($p_{\mathrm{dev}}$) is strictly positive. (B) Scenario for failed technology development: the probability that the technology transitions from *idle* to *ready* when investing in R&D ($p_{\mathrm{dev}}$) is null.

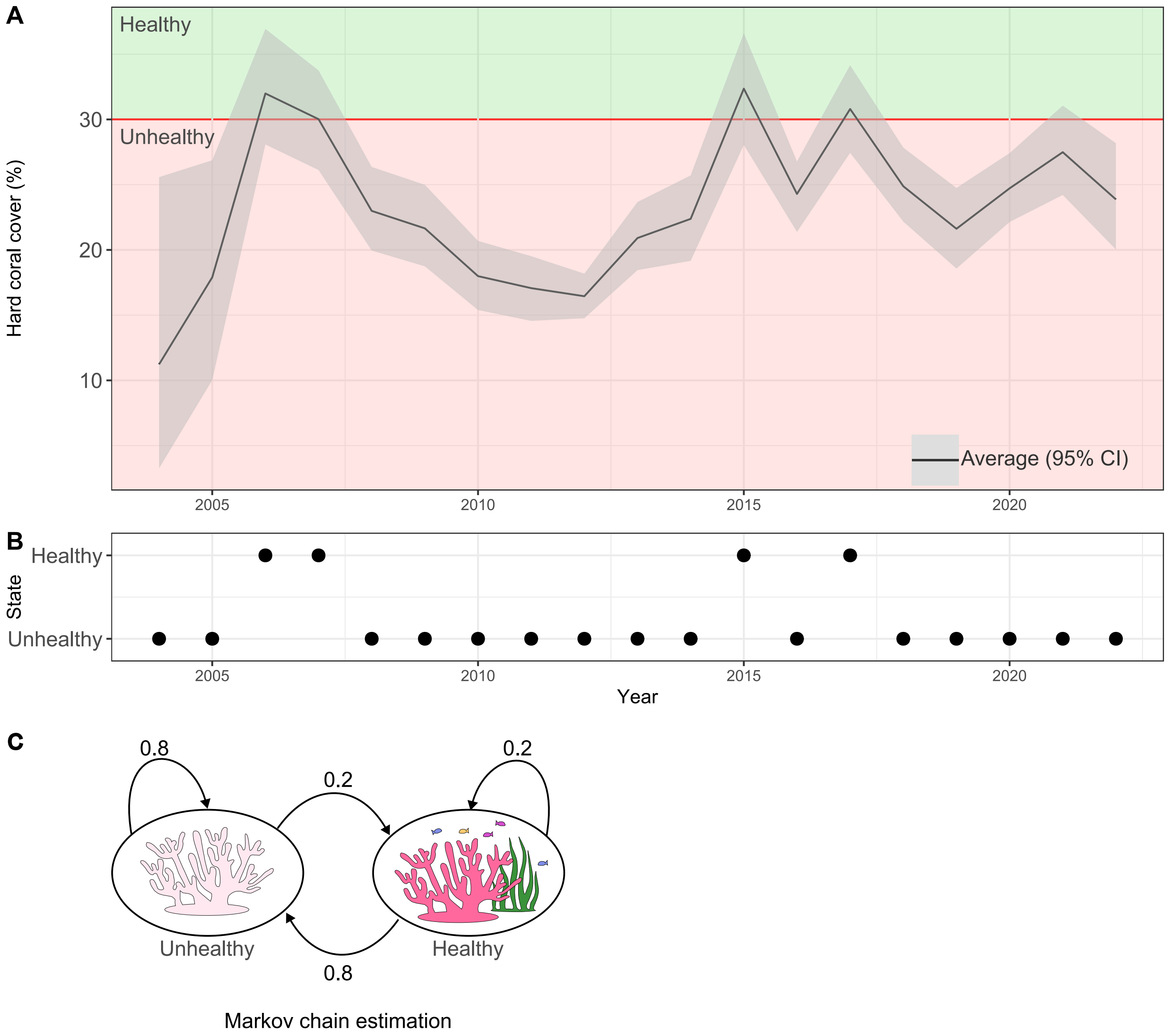


Figure S2: **A.** Time series representation of hard coral cover for the Central and Southern Great Barrier Reef (source: ReefCloud [1]). **B.** State classification as healthy (hard coral cover above 30%) or unhealthy (hard coral cover under 30 %). **C.** Markov chain estimation for the action business as usual.

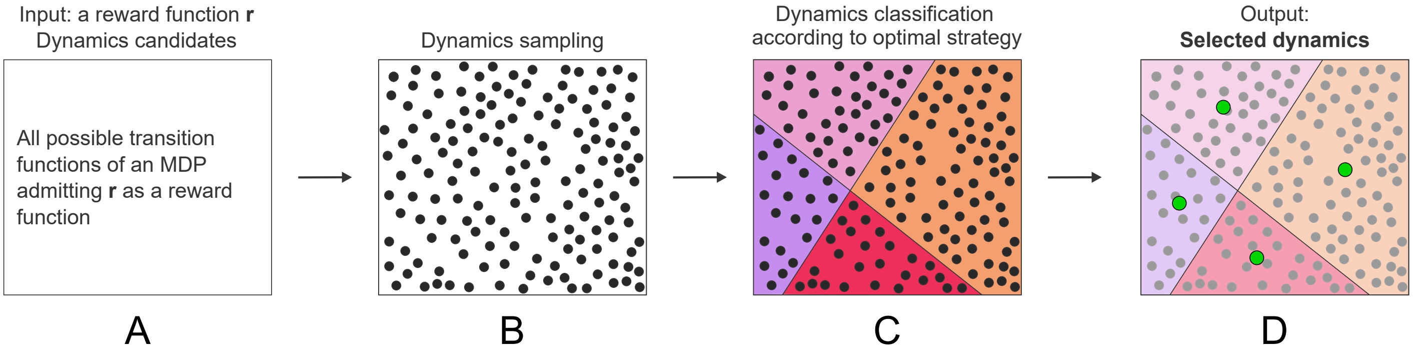


Figure S3: Steps of the 2-state n-action adaptive management solver [9]. This algorithm inputs a reward function of an MDP with up to 2 states and a finite number of actions, and returns a finite small and robust set of possible scenarios for adaptive management under model uncertainty. (A) First, the algorithm considers all the possible transition functions of an MDP admitting as reward function the input $r$. Then, the algorithm samples the set of candidates (B), and classifies each sampled dynamics depending on its optimal strategy: each color represents a different strategy (C). For each strategy, the algorithm averages the sampled dynamics: these are the outputs of the algorithm (D).

**A.** Universal model when business as usual is the most cost-efficient strategy ($m_1$)

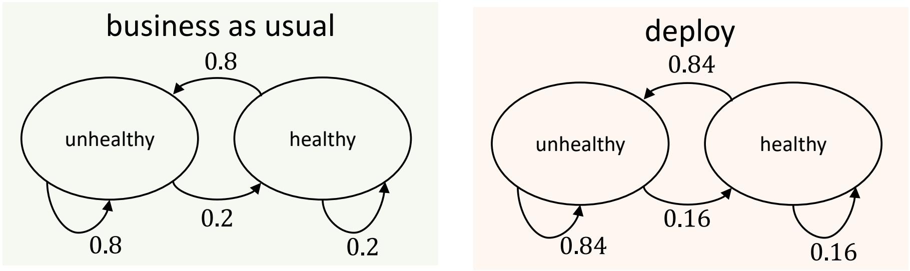


**B.** Universal model when business as usual is the more cost-efficient action when unhealthy and deploying new technology when healthy ($m_2$)

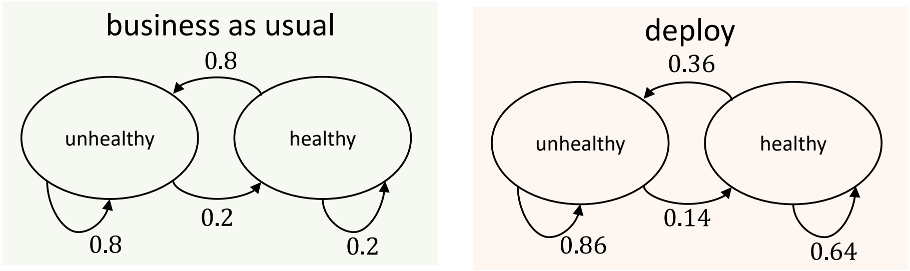


**C.** Universal model when business as usual is the more cost-efficient action when healthy and deploying new technology when unhealthy ($m_3$)

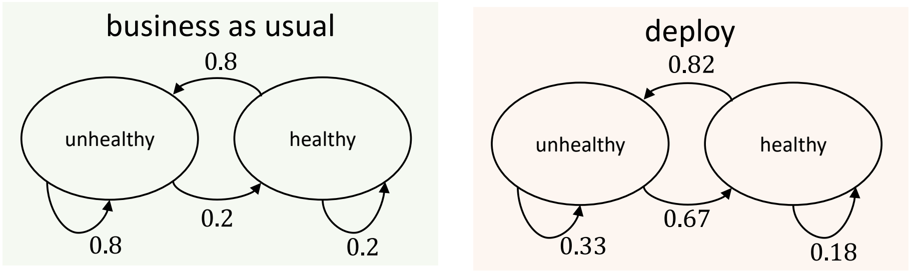


**D.** Universal model when deploying new technology is the most cost-effective strategy ($m_4$)

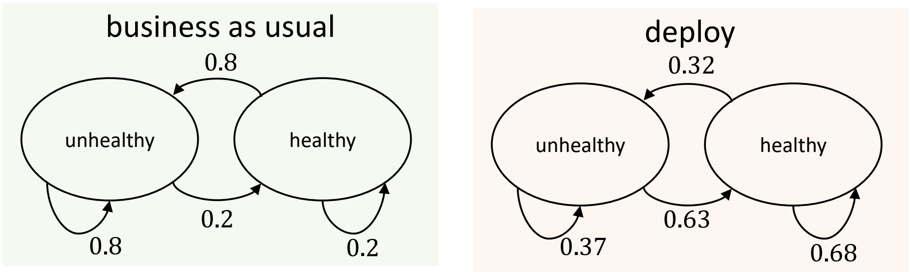


Figure S4: Universal models representatives of each possible optimal policy for the Great Barrier Reef case study. **A-D** represent 4 possible scenarios for technology deployment. The parameters are calculated with the 2-state n-action universal adaptive management solver (Section B.3). The parameters describing the ecosystem dynamics for the action *buisiness-as-usual* are assumed fully known (see Section B.1) and are the same across all scenarios.

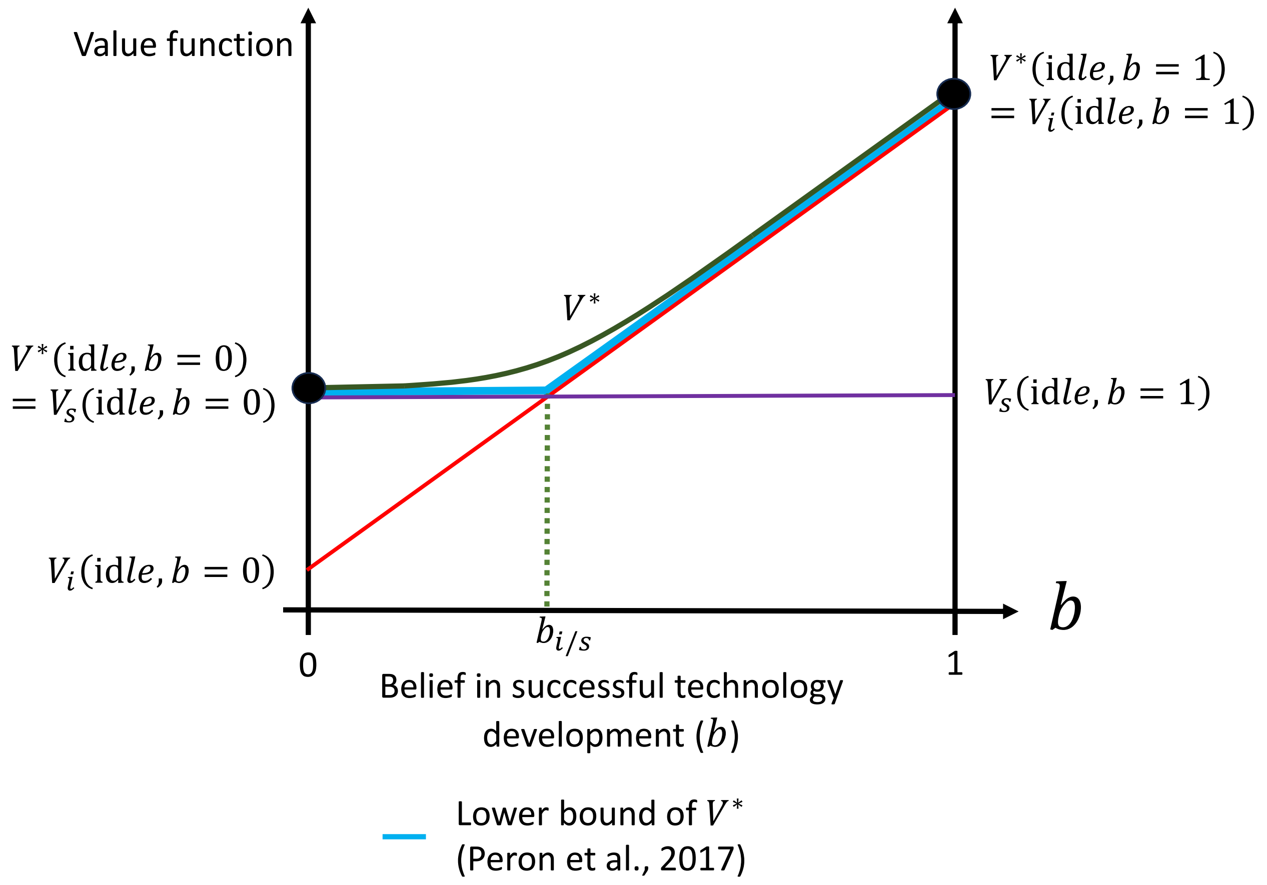


Figure S5: Approximation of belief state where optimal policy switches from invest in technology development to surrender. We approximate this belief state building on the lower bound approximation of Peron et al. [10].

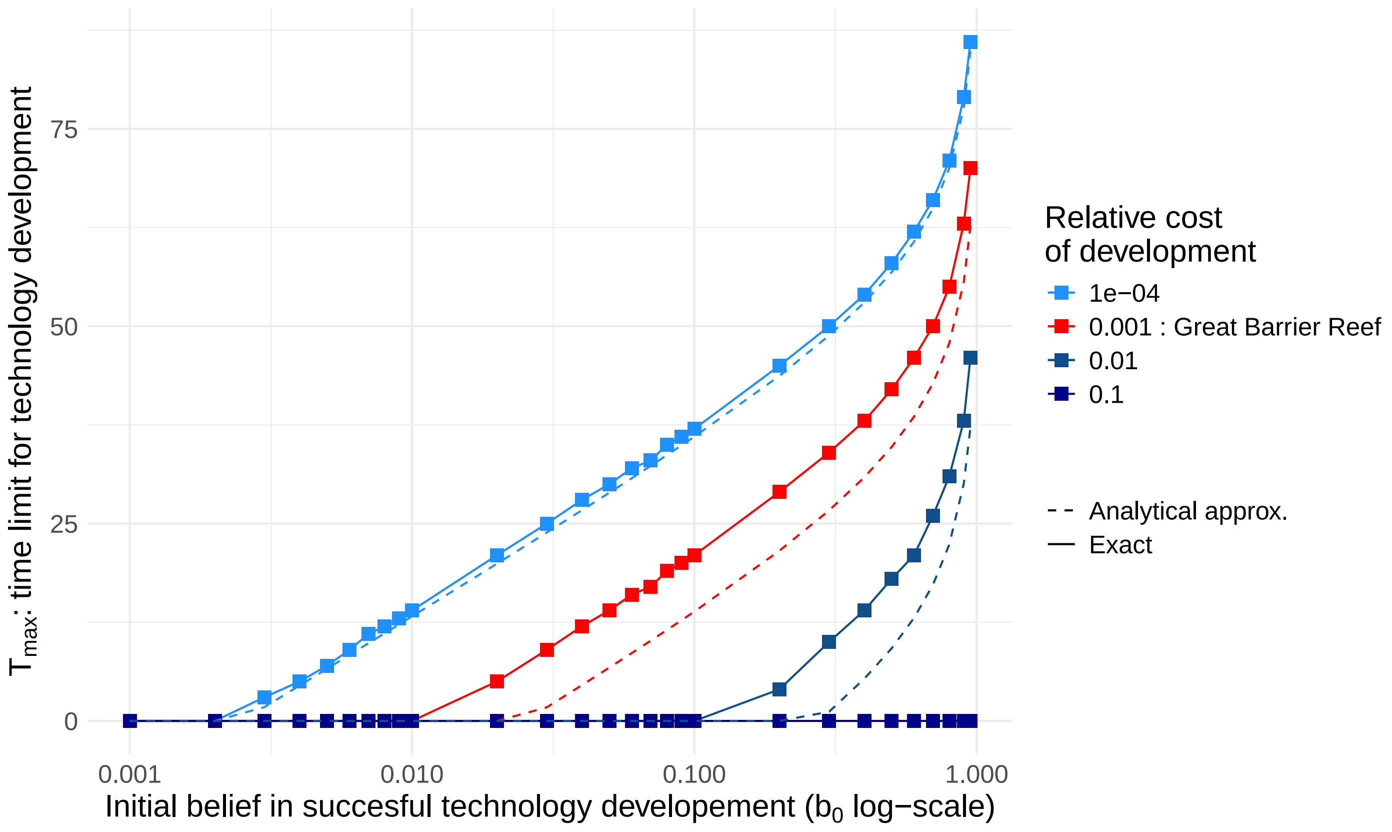


Figure S6: Maximum number of years investing in project development for different initial beliefs in R&D program feasibility, and different project development costs. Solid lines represent exact solution (POMDP) and dashed lines the approximate analytical solution. Here, $p_{\text{dev}} = 0.1$, $p_{\text{d}} = 0.8$, $p_{\text{r}} = 0.2$

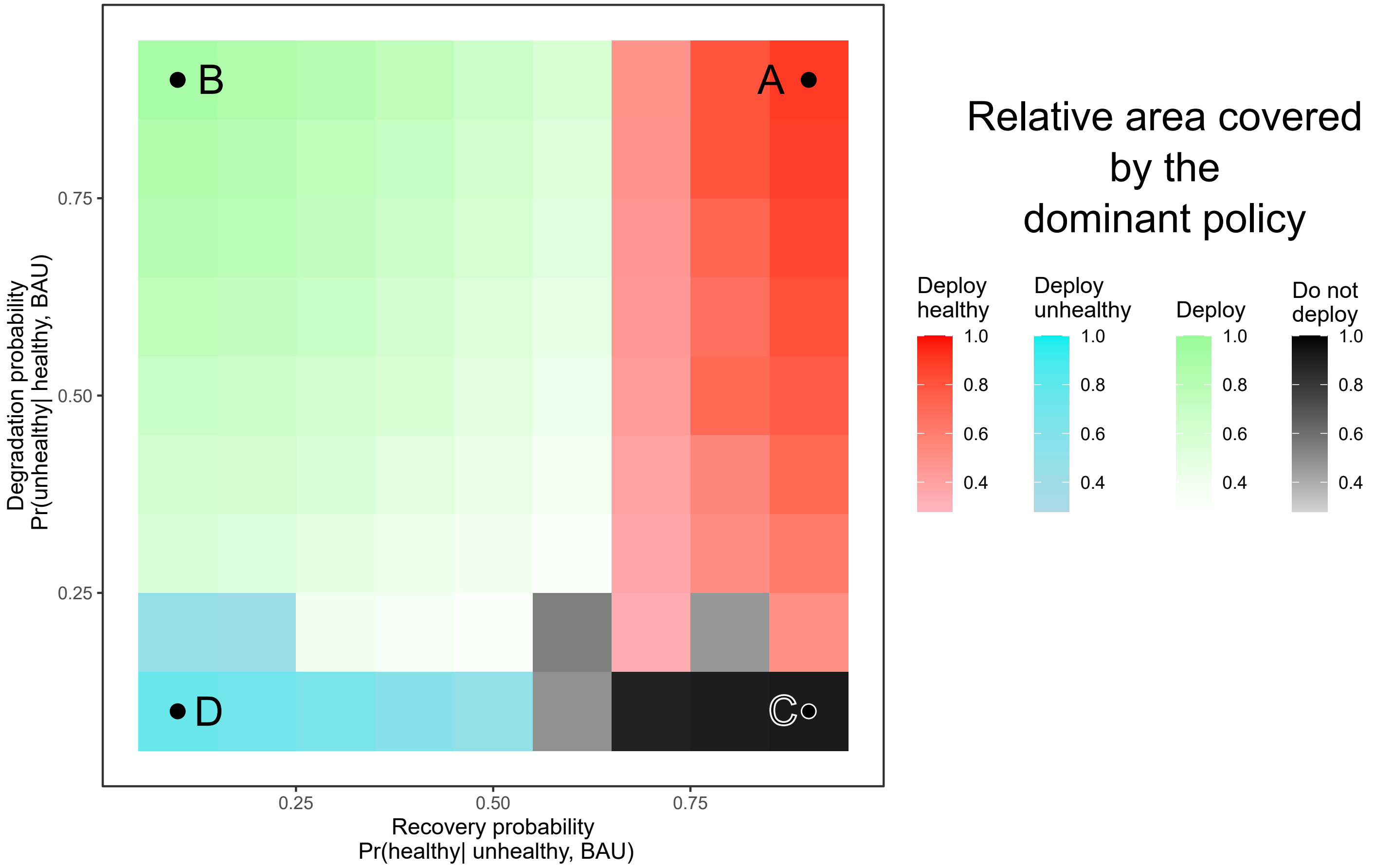


Figure S7: Summary of adaptive management strategies across possible ecosystem profiles. The x-axis represents the recovery probability of the ecosystem when the action *business-as-usual* (BAU) is implemented ($P(\text{healthy}|\text{unhealthy, BAU})$). The y-axis the degradation probability when the action *business-as-usual* (BAU) is implemented ($P(\text{unhealthy}|\text{healthy, BAU})$). See Section B.1. Here, A, B, C, D represent the ecosystem profiles studied in the main document and corresponding to different degradation-recovery profiles (see Figure 4). Each color depicts the dominant strategy for technology deployment (see Section B.2), and the shade of each color represents the relative importance of this dominant strategy compared to the other ones. As shown in the main document, the dominant strategy can be used as a proxy to determine the importance of the initial beliefs in the capacity of the technology for restoration ($\beta_0^{\mathrm{r}}$), and degradation prevention ($\beta_0^{\mathrm{p}}$).

Table S1: Summary of notations with values of parameters for the Great Barrier Reef case study. Costs and values of the reward function are relative to maximum possible benefits ($B_{\mathrm{h}}$).

| **Costs** | | | |
|---|---|---|---|
| Description | Notation | Cost (AUD) | Relative cost |
| Developing technology | $C_{\mathrm{dev}}$ | 5 Million | 0.001 |
| Deploying technology | $C_{\mathrm{deploy}}$ | 10 million | 0.002 |
| Business as usual | $C_{\text{BAU}}$ | 0 | 0 |
| **Benefits** | | | |
| Description | Notation | Value (AUD) | Relative value |
| Yearly ecosystem services produced by the reef when the reef is healthy | $B_{\mathrm{h}}$ | 6.4 Billion | 1 |
| Yearly ecosystem services produced by the reef when the reef is unhealthy | $B_{\mathrm{u}}$ | 4.3 Billion | 0.67 |
| Yearly expected ecosystem services generated by the reef under current management | $R_{\mathrm{BAU}}$ | - | - |
| Yearly expected ecosystem services generated by the reef under technology deployment with adaptive management | $R_{\mathrm{AM}}$ | - | - |
| **Dynamics parameters** | | | |
| Description | Notation | Value | |
| Probability of successful development if technology development scenario is *success* $P(\text{ready}\|\text{idle, invest in development, success})$ | $p_{\mathrm{dev}}$ | 0.1 | |
| Probability of successful development if technology development scenario is *failure* $P(\text{ready}\|\text{idle, invest in development, failure})$ | - | 0 | |
| Probability of degradation $P(\text{unhealthy}\|\text{healthy, BAU})$ | $p_{\mathrm{d}}$ | 0.8 | |
| Difficulty of recovery $P(\text{unhealthy}\|\text{unhealthy, BAU})$ | $p_{\mathrm{r}}$ | 0.8 | |
| **Initial beliefs** | | | |
| Description | Notation | Value | |
| Initial belief that the project will be successfully developed | $b_0$ | 0.5 | |
| Initial belief technology will be more cost-effective than business as usual when the reef is healthy (for preventing degradation) | $\beta^{\mathrm{p}}$ | 0.8 | |
| Initial belief technology will be more cost-effective than business as usual when the reef is unhealthy (for restoration) | $\beta^{\mathrm{r}}$ | 0.8 | |

Table S2: Reward achieved for the technology development POMDP, defined for each state and each action. Note here that the variable $R_{\mathrm{dep}}$ represents the expected benefits of deployment, and includes deployment costs.

| Development stage | surrender | invest in R&D or deploy |
|---|---|---|
| idle | $R_{\mathrm{BAU}}$ | $R_{\mathrm{BAU}} - C_{\mathrm{dev}}$ |
| ready | $R_{\mathrm{BAU}}$ | $R_{\mathrm{dep}}$ |

Table S3: Reward achieved for the technology deployment POMDP, defined for each state and each action.

| Ecosystem health | business as usual | deploy |
|---|---|---|
| unhealthy | $B_{\mathrm{u}}$ | $B_{\mathrm{u}} - C_{\mathrm{deploy}}$ |
| healthy | $B_{\mathrm{h}}$ | $B_{\mathrm{h}} - C_{\mathrm{deploy}}$ |

Table S4: Dynamic Programming approach to determine $V_i(\text{idle}, b = [1, 0])$ (value function of the technology development POMDP when the technology is idle and technology development scenario is *success*). At each time step, the value function is calculated as the sum of (a) the instant value of benefits and costs at the current state and (b) the weighted sum of benefits of the future time step for each possible state, which is discounted by $\gamma$. The later is $1 - p_{\text{dev}}$ the probability of the technology staying *idle* when investing in R&D multiplied by the future benefits of being in an *idle* state, plus $p_{\text{dev}}$ (probability of becoming *ready*) multiplied by $R_{\text{AM}}(\sum_{i=0}^{t} \gamma^i)$. We study the limit when $t \to +\infty$ to evaluate the long term benefits of the *invest in R&D* strategy. See Section C.1 for further details.

| Time | $V_i(\text{idle}, b = 1)$ | $V_i(\text{ready}, b = 1)$ |
|---|---|---|
| 0 | $R_{\text{BAU}}$ | $R_{\text{AM}}$ |
| 1 | $R_{\text{BAU}} - C_{\text{dev}} + (1 - p_{\text{dev}})\gamma R_{\text{BAU}} + p_{\text{dev}}\gamma R_{\text{AM}}$ | $R_{\text{AM}}(1 + \gamma)$ |
| 2 | $R_{\text{BAU}} - C_{\text{dev}} + (1 - p_{\text{dev}})\gamma[R_{\text{BAU}} - C_{\text{dev}} + (1 - p_{\text{dev}})\gamma R_{\text{BAU}} + p_{\text{dev}}\gamma R_{\text{AM}}] + p_{\text{dev}}\gamma R_{\text{AM}}(1 + \gamma)$ $= (R_{\text{BAU}} - C_{\text{dev}})(1 + (1 - p_{\text{dev}})\gamma) + (1 - p_{\text{dev}})^2\gamma^2 R_{\text{BAU}} + p_{\text{dev}}\gamma R_{\text{AM}}[(1 - p_{\text{dev}})\gamma + (1 + \gamma)]$ | $R_{\text{AM}}(1 + \gamma + \gamma^2)$ |
| ⋮ | | |
| $t$ | $(R_{\text{BAU}} - C_{\text{dev}})(\sum_{i=0}^{t}(1 - p_{\text{dev}})^i\gamma^i) + (1 - p_{\text{dev}})^t\gamma^t R_{\text{BAU}} + p_{\text{dev}}\gamma R_{\text{AM}}(\sum_{i=0}^{t-1}(1 - p_{\text{dev}})^i\gamma^i[\sum_{j=0}^{t-i-1} \gamma^j])$ | $R_{\text{AM}}(\sum_{i=0}^{t} \gamma^i)$ |
| ⋮ | | |
| $+\infty$ | $\frac{R_{\text{BAU}} - C_{\text{dev}} + p_{\text{dev}}\gamma W^*}{1 - (1 - p_{\text{dev}})\gamma}$ | $R_{\text{AM}}(\sum_{i=0}^{\infty} \gamma^i) = W^*$ |

Table S5: Data set of hard coral cover (%) for the Central and Southern Great Barrier Reef (source: ReefCloud [1]). The upper and lower bounds represent the 95% confidence intervals. This data set was used to generate Figure S2, and estimate the ecosystem dynamics under business-as-usual activities (see Section B.1).

| Upper bound | Average | Lower bound | year |
|---|---|---|---|
| 25.58 | 11.23 | 3.26 | 2004 |
| 26.87 | 17.9 | 10.02 | 2005 |
| 36.94 | 31.99 | 28.08 | 2006 |
| 33.76 | 30.02 | 26.12 | 2007 |
| 26.36 | 22.99 | 19.95 | 2008 |
| 24.99 | 21.65 | 18.74 | 2009 |
| 20.69 | 17.99 | 15.39 | 2010 |
| 19.52 | 17.07 | 14.56 | 2011 |
| 18.18 | 16.45 | 14.76 | 2012 |
| 23.67 | 20.91 | 18.45 | 2013 |
| 25.71 | 22.38 | 19.16 | 2014 |
| 36.62 | 32.36 | 28.05 | 2015 |
| 26.78 | 24.29 | 21.38 | 2016 |
| 34.16 | 30.8 | 27.44 | 2017 |
| 27.84 | 24.88 | 22.17 | 2018 |
| 24.75 | 21.62 | 18.57 | 2019 |
| 27.42 | 24.72 | 22.14 | 2020 |
| 31.06 | 27.49 | 24.21 | 2021 |
| 28.18 | 23.87 | 20.03 | 2022 |
